# The GENIUS Approach to Robust Mendelian Randomization Inference

**DOI:** 10.1101/193953

**Authors:** Eric J. Tchetgen Tchetgen, BaoLuo Sun, Stefan Walter

## Abstract

Mendelian randomization (MR) is a popular instrumental variable (IV) approach, in which one or several genetic markers serve as IVs that can be leveraged to recover under certain conditions, valid inferences about a given exposure-outcome causal association subject to unmeasured confounding. A key IV identification condition known as the exclusion restriction states that the IV has no direct effect on the outcome that is not mediated by the exposure in view. In MR studies, such an assumption requires an unrealistic level of knowledge and understanding of the mechanism by which the genetic markers causally affect the outcome, particularly when a large number of genetic variants are considered as IVs. As a result, possible violation of the exclusion restriction can seldom be ruled out in such MR studies, and if present, such violation can invalidate IV-based inferences even if unbeknownst to the analyst, confounding is either negligible or absent. To address this concern, we introduce a new class of IV estimators which are robust to violation of the exclusion restriction under a large collection of data generating mechanisms consistent with parametric models commonly assumed in the MR literature. Our approach which we have named “MR G-Estimation under No Interaction with Unmeasured Selection” (MR GENIUS) may in fact be viewed as a modification to Robins’ G-estimation approach that is robust to both additive unmeasured confounding and violation of the exclusion restriction assumption. We also establish that estimation with MR GENIUS may also be viewed as a robust generalization of the well-known Lewbel estimator for a triangular system of structural equations with endogeneity. Specifically, we show that unlike Lewbel estimation, MR GENIUS is under fairly weak conditions also robust to unmeasured confounding of the effects of the genetic IVs on both the exposure and the outcome, another possible violation of a key IV Identification condition. Furthermore, while Lewbel estimation involves specification of linear models both for the outcome and the exposure, MR GENIUS generally does not require specification of a structural model for the direct effect of invalid IVs on the outcome, therefore allowing the latter model to be unrestricted. Finally, unlike Lewbel estimation, MR GENIUS is shown to equally apply for binary, discrete or continuous exposure and outcome variables and can be used under prospective sampling, or retrospective sampling such as in a case-control study, as well as for right censored time-to-event outcomes under an additive hazards model.

## 1 Introduction

Mendelian randomization (MR) is an instrumental variable approach with growing popularity in epidemiology studies. In MR, one aims to establish a causal association between a given exposure and an outcome of interest in the presence of possible unmeasured confounding, by leveraging one or more genetic markers defining the IV (Davey Smith and Ebrahim, 2003, 2004, Lawlor et al, 2008). In order to be valid IVs, the genetic markers must satisfy the following key conditions:

a. They must be associated with the exposure.
b. They must be independent of any unmeasured confounder of the exposure-outcome relationship.
c. There must be no direct effect of a genetic marker on the outcome that is not fully mediated by the exposure in view.

The last assumption (c) known as the exclusion restriction is rarely credible in the context of MR as it requires complete understanding of the biological mechanism by which each marker influences the outcome. Such a priori knowledge may be unrealistic in practice due to the possible existence of unknown pleitropic effects of the markers (Little and Khoury, 2003; Davey Smith and Ebrahim 2003, 2004, Lawlor et al 2008). Violation of assumption (b) can also occur due to linkage disequilibrium or population stratification (Lawlor et al, 2008). Possible violation or near violation of assumption (a) known as the weak instrumental variable problem also poses an important challenge in MR as individual genetic effects on phenotypes can be fairly weak.

There has been tremendous interest in the development of formal statistical methods to detect and account for violation of IV assumptions (a)-(c), primarily in a multiple-IV setting in which standard linear models for outcome and exposure are assumed. The literature addressing violation of assumption (a) is arguably the most developed and extends to possible nonlinear models under a generalized methods of moments framework; some recent papers on this topic include Staiger and Stock (1997), Stock and Wright (2000), Stock and Yogo (2002), Chao and Swanson (2005). Methodology to address violations of (b) or (c) is far less developed, and constitutes the central focus of this paper. Three strands of work stand out in recent literature concerning violation of either of these assumptions. In the first strand, Kang et al (2016) developed a penalized regression approach that can under certain conditions recover valid inferences about the causal effect of interest provided fewer than fifty percent of genetic markers are invalid IVs; also see Windmeijer et al (2016) for improvements on the penalized approach of Kang et al (2016), including a proposal for standard error estimation which was not provided in Kang et al (2016). In an alternative approach, Han (2008) established that the median of multiple estimators of the effect of exposure obtained using one instrument at the time is a consistent estimator also assuming fewer than fifty percent of IVs are invalid and that IVs cannot have direct effects on the outcome unless the IVs are uncorrelated. Bowden et al (2016) explore closely related weighted median methodology. In a second strand of work, Guo et al (2017) proposed two stage hard thresholding (TSHT) with voting, which is able to recover a consistent causal effect estimator under linear models for the outcome and exposure, and a certain plurality condition which can be considerably weaker than the fifty percent rule (also known as majority rule). The plurality condition is defined in terms of regression parameters encoding the association of each invalid IV with the outcome and that encoding the association of the corresponding IV with the exposure. The condition effectively requires that the number of valid IVs is greater than the largest number of invalid IVs with equal ratio of the above regression coefficients. Furthermore, they provide a simple construction for 95% confidence intervals to obtain inferences about the exposure effect which are guaranteed to have correct coverage under the plurality condition. Importantly, in these first two strands of work, a candidate IV may be invalid either because it violates the exclusion restriction, or because it shares an unmeasured common cause with the outcome, i.e. either (b) or (c) fails. Both the penalized approach and the median estimator may be inconsistent if 50% or more candidate IVs turnout to be invalid, while TSHT may be inconsistent if the plurality rule fails. For instance, it is clear that neither approach can recover valid inferences if all IVs violate either assumption (b) or (c). In order to remedy this difficulty, in a third strand of work, Kolesar et al (2011) considered the possibility of identifying the exposure causal effect when all IVs violate the exclusion restriction (c), provided the effects of the IVs on the exposure are asymptotically orthogonal to their direct effects on the outcome as the number of IVs tends to infinity. A closely related meta-analytic version of their approach known as MR-Egger has recently emerged in the epidemiology literature (Bowden et al, 2015); they referred to the orthogonality condition as the instrument strength independent of direct effect (InSIDE) assumption. As pointed out by Kang et al (2016), the orthogonality condition on which these approaches rely may be hard to justify in MR settings as it potentially restricts unknown pleitropic effects of the genetic markers often with little to no biological basis. A notable feature of aforementioned methods is that they are primarely tailored to a multiple-IV setting, in fact methods such as MR-Egger are consistent only under an asymptotic theory in which the number of IVs goes to infinity, together with sample size. It is also important to note that because confidence intervals for the causal effect of the exposure obtained by Windmeijer et al (2015) and Guo et al (2017) rely on a consistent model selection procedure, such confidence intervals fail to be uniformly valid over the entire model space (Guo et al, 2017, Leeb and Potscher, 2008).

Because in practice, it is not possible to ensure that fewer than fifty percent of candidate IVs are invalid or that the plurality condition holds, nor is it practically possible to enforce the orthogonality condition of Kolesar et al (2011), an important goal of MR research aims to develop alternative methods of estimation and inference that are fully robust to possible violation of IV assumptions without relying on majority, plurality or orthogonality conditions. In this paper, a class of estimators fulfilling this desideratum is proposed, which unlike the aforementioned robust methods equally applies whether one has observed a single or many candidate IVs.

In Section 2, we introduce notation used throughout. We also provide a formal definition of the IV model for which we describe previously proposed sufficient conditions in the canonical case of binary exposure and IV, for nonparametric identification of the exposure average causal effect in terms of the so-called Wald estimand (Wang and Tchetgen Tchetgen, 2017). In Section 3, we present our first result which provides an alternative identification formula for the average causal effect in the IV context, which unlike the Wald estimand, is robust to violation of the exclusion restriction (c) under a large collection of possible data generating mechanisms that assume both (i) no additive interaction between the exposure, an unmeasured confounder and the candidate IV in a mean model for the outcome; and (ii) no additive interaction between the candidate IV and an unmeasured confounder in a mean model for the exposure. Conditions similar to assumptions (i) and (ii) are fairly common in MR and other IV literature. For instance, Kolesar et al (2011), Kang et al (2016) and Bowden et al (2015) rely on analogous assumptions. In Section 3, we establish that the proposed approach readily accounts for continuous exposure. We establish that our approach which we call”MR G-Estimation under No Interaction with Unmeasured Selection” (MR GENIUS) may in fact be viewed as a modification to Robins’ G-estimation approach (Robins, 1997) which we have made robust to both additive unmeasured confounding and violation of the exclusion restriction assumption. Identification with MR GENIUS relies primarily on an assumption that the conditional variance of the exposure given the candidate IVs is heteroscedastic with respect to the candidate IVs, an assumption which generally holds for binary or discrete exposure except at certain exceptional data generating mechanisms. In case of continuous exposure and outcome, this assumption is closely related to Lewbel’s recent proposal to leverage heteroscedasticity for identification and estimation in endogenous regression models (Lewbel, 2012). In this case, MR GENIUS and Lewbel estimation are quite similar, although unlike estimation with the Lewbel approach, estimation with MR GENIUS avoids specification of a model for the direct effect of invalid IVs with the outcome, therefore allowing the latter to remain unrestricted. In Section 4, we describe conditions under which MR GENIUS is also robust to unmeasured confounding of the effects of the genetic IVs on both the exposure and the outcome, a violation of assumption (b) which is also not appropriately accounted for by Lewbel regression which assumes that candidate IVs are independent of unmeasured confounders. As we further establish inSection 5, MR GENIUS can easily incorporate multiple IVs in a generalized methods of moments (GMM) approach. An important feature of multiple IV MR GENIUS is that the correlation structure for the IVs can essentially remain unrestricted without necessarily affecting identification, this is in contrast with Bowden et al (2015) who require uncorrelated IVs and Kang et al (2016) who likewise require IV correlation structure to be somewhat restricted (Windmeijer et al, 2015). Section 5 also extends the proposed approach to target a multiplicative average causal effect, and establishes that in the case of binary outcome, the approach is equally valid under either prospective or retrospective sampling designs. Therefore, MR GENIUS can also be viewed as further generalizing Lewbel’s estimator to these important settings. In Section 5, we also briefly extend MR GENIUS to the context of a right censored time-to-event endpoint under a structural additive hazards model, therefore further robustifying the recent semiparametric IV estimator of Martinussen et al (2017) against possible violation of the exclusion restriction assumption. In Section 6, we evaluate the proposed methods and compare them to a number of previous MR methods in extensive simulation studies. In Section 7 we illustrate the methods in an MR analysis of the effect of diabetes on memory in the Health and Retirement Study. Section 8 offers some concluding remarks.

## 2 Notation and definitions

Suppose that one has observed *n* i.i.d. realizations of a vector (*A, G, Y*) where *A* is an exposure, *G* the candidate IV and *Y* is the outcome. Let *U* denote an unmeasured confounder (possibly multivariate) of the effect of *A* on *Y. G* is said to be a valid instrumental variable provided it fulfills the following three conditions:

**Assumption 1.** IV relevance: *G /⊥ ⊥ A|U*;

**Assumption 2.** IV independence: *G⊥ ⊥ U*;

**Assumption 3.** Exclusion restriction: *G⊥ ⊥ Y |A, U.*

The first condition ensures that the IV is a correlate of the exposure even after conditioning on *U.* The second condition states that the IV is independent of all unmeasured confounders of the exposure-outcome association, while the third condition formalizes the assumption of no direct effect of *G* on *Y* not mediated by *A* (assuming Assumption 2 holds). The causal diagram in Figure 1 encodes these three assumptions and therefore provides a graphical representation of the IV model. It is well known that while a valid IV satisfying assumptions 1-3, i.e. the causal diagram in Figure 1, suffices to obtain a valid statistical test of the sharp null hypothesis of no individual causal effect, the population average causal effect is itself not point identified with a valid IV without an additional assumption. Consider the following condition:

**Assumption 4.**

**(4a)** There is no additive *A -* (*U, G*) interaction in model for *E* (*Y |A, G, U*)

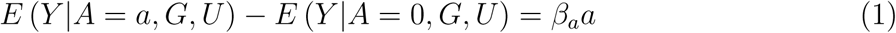

and no additive *G -* (*U*) interaction in model for *E* (*Y |A, G, U*)

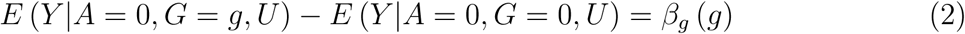

for an unknown function *β*_*g*_ (*·*) that satisfies *β*_*g*_ (0) = 0

**(4b)** There is no additive *G - U* interaction in model for *E* (*A|G, U*)

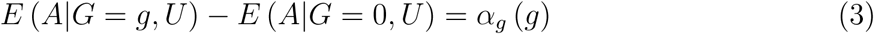

for an unknown function *α*_*g*_ (*·*) that satisfies *α*_*g*_ (0) = 0.

Clearly the condition does not require *G* to be a valid IV. Equation (1) implies that the average causal effect of *A* on *Y* conditional on *U* and *G* does not depend on *U* and *G* on the additive scale, i.e. the additive causal effect of *A* on *Y* is not modified by either *U* or *G*. Likewise, equation (2) additionally states that the additive average effect of *G* on *Y* is not modified by *U.* These restrictions imply the following additive models:

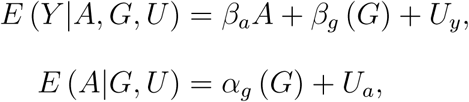

where *U*_*y*_ = *β*_*u*_ (*U*) and *U*_*a*_ = *α*_*u*_(*U*) for functions *β*_*u*_ (*·*) and *α*_*u*_(*·*) only restricted by natural features of the model, e.g. such that the outcome and exposure means are bounded between zero and one in the binary case. If *G* is a valid IV, then *E* (*Y |A, G, U*) = *E* (*Y |A, U*) does not depend on *G* by the exclusion restriction implying that *G* neither interacts with *U* nor with *A* in the model for *E* (*Y |A, G, U*), so that assumption 4.a. reduces to the assumption of no *U – A* interaction in the model for *E* (*Y |A, G, U*). In case of a valid binary IV and binary exposure, Wang and Tchetgen Tchetgen (2017) recently established that the average causal effect *β*_*a*_ is nonparametrically identified by the so-called Wald estimand

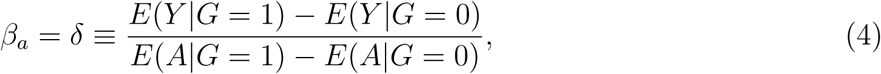

if either of 4.a. or 4.b holds but not necessarily both conditions hold. Note that the models for *E*(*Y |A, G, U*) and *E*(*A|G, U*) considered by Bowden et al (2015) satisfy assumptions 4.a. and 4.b. with *β*_*g*_ (*·*) and *α*_*g*_ (*·*) linear functions, while Kang et al (2016) specified models implied by these two restrictions. Below, unless stated otherwise, assume *A* and *G* are both binary.

**Figure 1.**
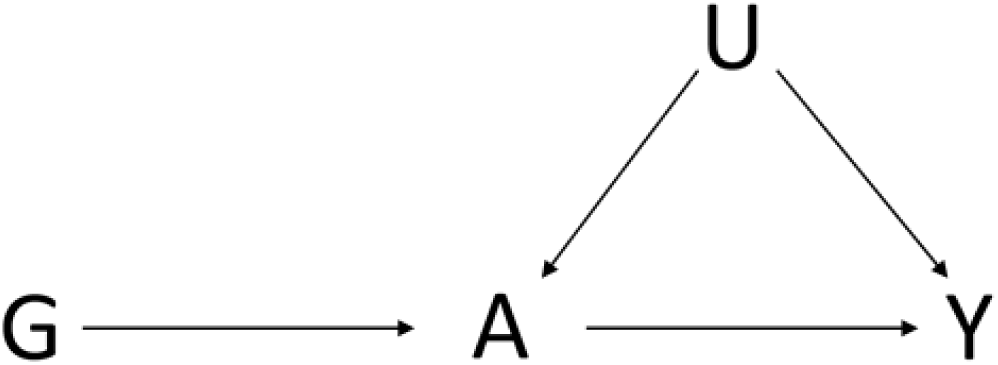
Directed acyclic graph depicting a valid instrument *G* which satisfies assumptions 1-3.

## 3 Identification under violation of exclusion restriction

Next, suppose that as encoded in the diagram given in Figure 2, the exclusion restriction assumption 3 does not necessarily hold, then the Wald estimand *δ ≠ β*_*a*_ will generally fail to equal the average causal effect of *A* on *Y,* even if assumptions 1, 2, and 4 hold. The following result provides an alternative identifying formula which may be used instead of the Wald estimand to identify the causal effect under these conditions.

### Lemma 1

*Suppose that Assumptions 1, 2 and 4 hold, then β*_*a*_ = *µ, where*

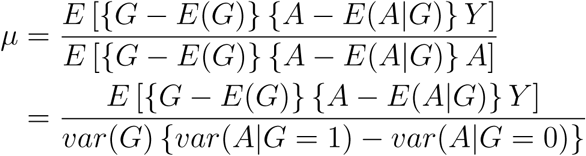

*provided that*

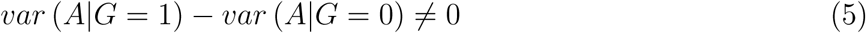

**Proof.** Below we make use of the fact that under our assumptions *E {A - E*(*A|G*)*|G, U}* = *α*_*u*_(*U*) *- E*(*α*_*u*_(*U*)). Consider

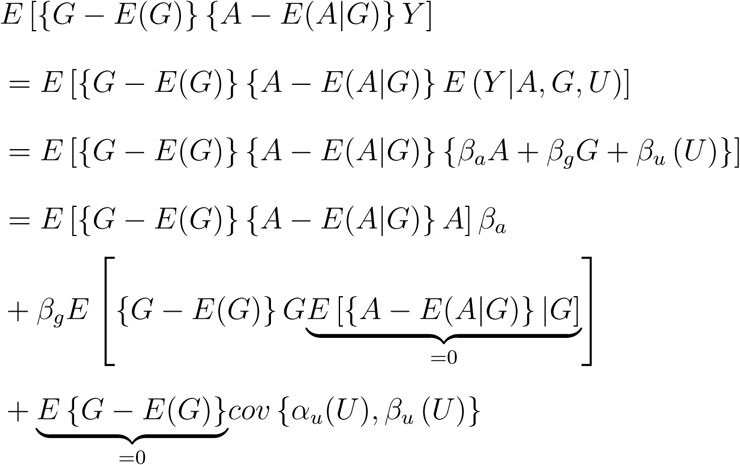

Therefore,

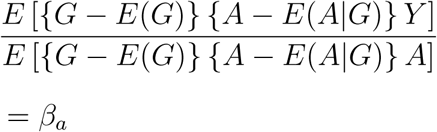

provided that *E* [{*G - E*(*G*)} {*A - E*(*A|G*)} *A*] *≠* 0, which holds under (5).

Lemma 1 provides an explicit identifying expression for the average causal effect *β*_*a*_ of *A* on *Y* in the presence of additive confounding, which leverages a candidate IV *G* that may or may not satisfy the exclusion restriction. In order for *µ* to be well defined, we require a slight strengthening of the IV relevance assumption 1,i.e. that *var*(*A|G*) must depend on *G*. It is key to note that this assumption is empirically testable, and will typically hold for binary *A*, except at certain exceptional laws. To illustrate, let *π*(*g*) = Pr(*A* = 1*|G* = *g*) and suppose that assumptions 1, 2 and 4 hold, however *π* (1) = 1*-π* (0), in which case (5) fails because *var*(*A|G* = *g*) = *π*(*g*) (1 *- π*(*g*)) = *π*(1) (1 *- π*(1)) = *π*(0) (1 *- π*(0)) does not depend on *g* and therefore the identifying expression given in the Lemma does not apply despite the candidate IV satisfying IV relevance assumption 1, i.e. *π* (1) *≠ π* (0). Below, we extend Lemma 1 to allow for possible violation of both assumptions 2 and 3.

The lemma motivates the following MR estimator, which is guaranteed to be consistent under assumptions 2, 4 and equation (5) irrespective of whether or not assumption 3 also holds:

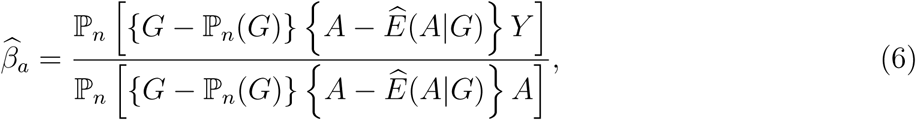

where 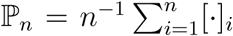 and Ê(*A|G* = *g*) = ℙ_*n*_ [*A*_*i*_1 (*G*_*i*_ = *g*)] */* ℙ_*n*_ [1 (*G*_*i*_ = *g*)]. This estimator is the simplest instance of MR GENIUS estimation. The asymptotic distribution of the estimator is described in Appendix A2.

**Figure 2.**
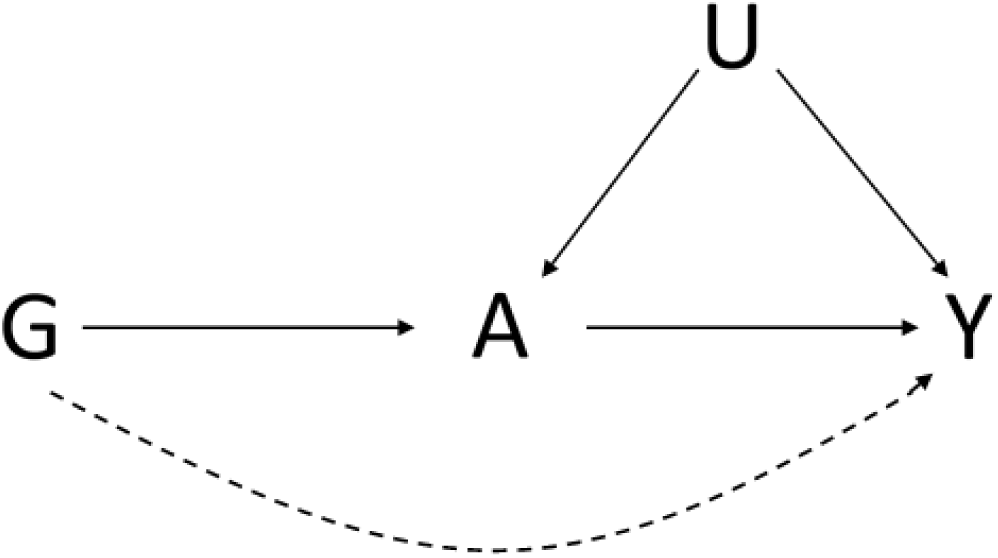
Directed acyclic graph depicting the situation in which exclusion restriction (assumption 3) does not necessarily hold. The dashed line indicates possible direct effect of *G* on outcome *Y*.

### Continuous exposure

Suppose now that *A* is continuous, then, it is straightforward to verify that Lemma 1 continues to hold as its proof does not depend on *A* being binary. Note that for continuous *A*, Assumption (1) resticts the effect of *A* on *Y* to be linear and condition (5) implies that the conditional density of *ε*_*A*_ = *A- E*(*A|G*) must be heteroscedastic, i.e. 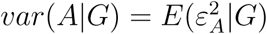 depends on *G.* As mentioned in the introduction, Lewbel (2012) obtains a closely related identification result to Lemma 1 under a triangular system of linear structural equation models; see Theorem 1 on Page 70 of Lewbel (2012). In addition to establishing the result for binary *A* in Lemma 1 without specification of a triangular system of linear equations, below we generalize this identification result in several important directions particularly relevant to MR studies.

We note that while *var*(*A|G*) will generally depend on *G* for binary or discrete *A* (except perhaps at exceptional data generating mechanisms such as the one described in the previous Section), this may not always be the case for continuous *A.* However in this case, the assumption can be motivated under an underlying model for *A* with latent heterogeneity in the effect of *G* on *A.* Specifically, suppose that

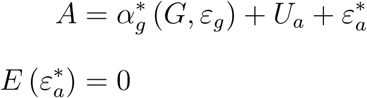

where *ε*_*g*_ and 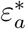 are unobserved random disturbances independent of (*G, U*); the disturbance *ε*_*g*_ may be viewed as unobserved genetic or environmental factors independent of *G*, that may however interact with *G* to induce additive effect heterogeneity of G-A associations, e.g. 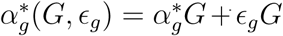. Then, one can verify that the model in the above display implies that *A* = *α*_*g*_ (*G*) + *ε*_*a*_ where 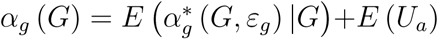 and 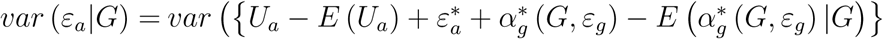 which clearly depends on *G*, provided 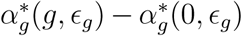 depends on ϵ_*g*_ for a value of *g*, therefore implying condition (5). A model for exposure which incorporates latent heterogeneity in the effects of *G* is quite natural in the MR context because such a model is widely considered a leading contestant to explain the mystery of missing heritability (Manolio et al, 2009).

## 4 Identification under violation of IV Independence

In this Section, we aim to relax the IV independence Assumption 2., by allowing for dependence between *U* and *G* as displayed in Figure 3. Therefore, we will consider replacing Assumption 2 with the following weaker condition:

**Assumption 2*.** Homoscedastic confounding: *cov* (*U*_*y*_*, U*_*a*_*|G*) = *ρ* does not depend on *G.*

To illustrate Assumption 2* it is instructive to consider the following submodels of (1) and (2): *U*_*y*_ = *β*_0_ + *β*_*u*_*U* and *U*_*a*_ = *α*_0_ + *α*_*a*_*U,* such that *E*(*Y |A, U, G*) and *E*(*A|G, U*) are both linear in *U*; then Assumption 2* implies *var*(*U|G*) = *ρ/* (*β*_*u*_*α*_*a*_), i.e. the unmeasured confounder *U* has homoscedastic variance. Under Assumption 2*, *E*(*U|G*) is left unrestricted therefore assumption 2 may not hold. We have the following result:

### Lemma 2

*Suppose that Assumptions 1, 2*,4 hold, then β*_*a*_ = *µ provided that condition* (5) *holds.*

**Proof.** Proceeding as in the proof of Lemma 1,

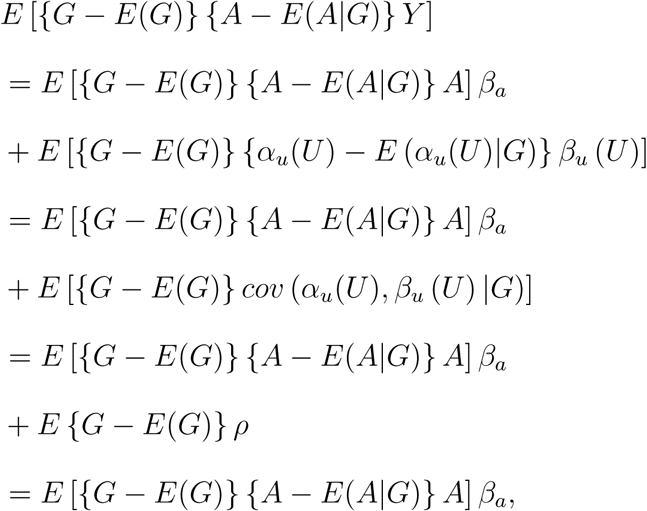

proving the result.

Lemma 2 implies that under Assumptions 1, 2*, 4 and condition (5) *, 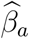* continues to be consistent even if *U /⊥ ⊥ G.*

As previously mentioned, MR GENIUS may be viewed as a special case of G-estimation (Robins, 1997). In fact, under assumption 4.a.1 and the additional assumption of no unobserved confounding given *G*, i.e. if either *U⊥ ⊥ A|G* or *U⊥ ⊥ Y |A, G,* the G-estimator 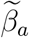 which solves an estimating equation of the form:

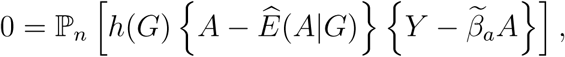

is consistent and asymptotically normal for any user-specified function *h* (*·*) (up to regularity conditions).

It is straightforward to verify that the MR GENIUS estimator (6) solves the estimating equation:

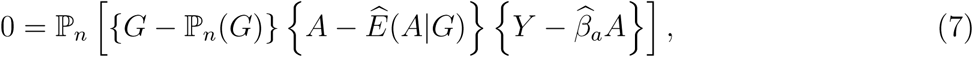

therefore formally establishing an equivalence between MR GENIUS and g-estimation for the choice *h*(*G*) = *G - E*(*G*). Remarkably, as we have established above, this specific choice of *h* renders g-estimation robust to unmeasured confounding under certain no-additive interactions conditions with unmeasured factors used in selecting exposure levels, therefore motivating the choice of acronym for the proposed approach.

**Figure 3.**
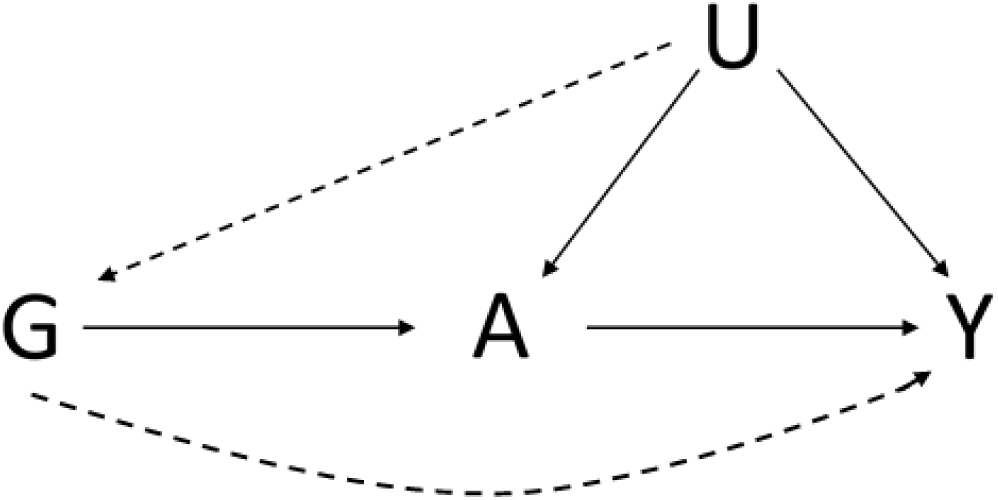
Directed acyclic graph depicting the situation in which IV independence (assumption 2) and exclusion restriction (assumption 3) do not necessarily hold. The dashed lines indicate possible direct effects of *U* on *G*, and of *G* on *Y*.

## 5 Generalizations

### 5.1 Multiplicative exposure model

A multiplicative exposure model may also be used for count or binary exposure under the following assumption:

**(4.b*)** There is no multiplicative *G - U* interaction in model for *E* (*A|G, U*)

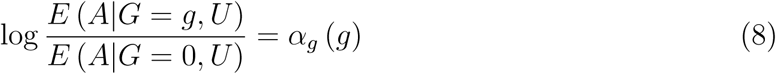

for an unknown function *α*_*g*_ (*·*) that satisfies *α*_*g*_ (0) = 0.

MR GENIUS can be adapted to this setting according to the following result. Let

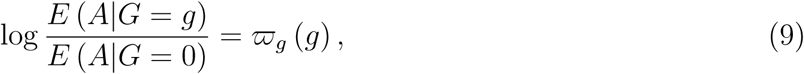

and *U*_*a*_ = *E* (*A|G* = 0*, U*).

#### Lemma 5

*Suppose that Assumptions 1, 2, 4.a and 4.b* hold, then*

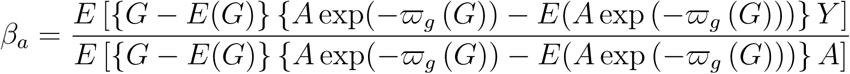

*provided that var* (*A|g*) */var* (*A|g* = 0) *≠* exp (*-ϖ*_*g*_ (*g*)) *for at least one value of g.*

**Proof.** The proof follows upon noting that under our assumptions,

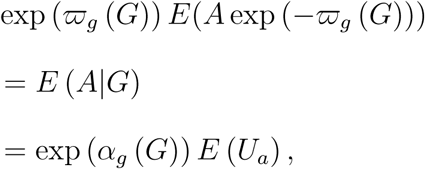

and

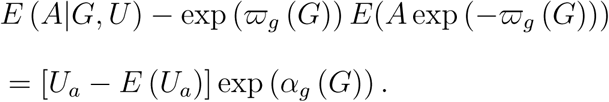

Therefore

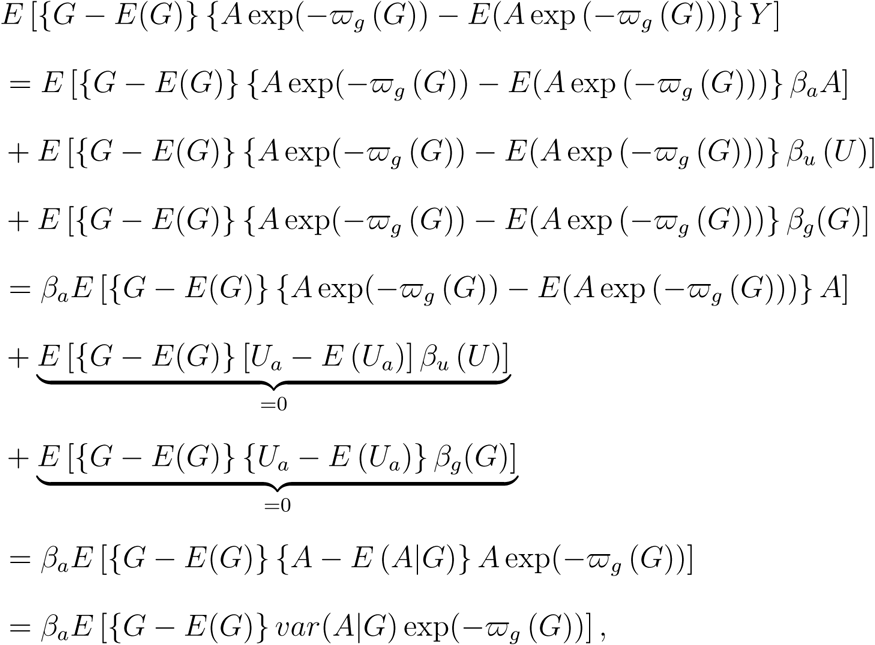

where we used the fact that under Assumption 2., *-ϖ*_*g*_ (*g*) = *α*_*g*_ (*g*), therefore proving identification provided that *var*(*A|G*) exp(*-ϖ*_*g*_ (*G*)) is a function of *G,*which holds as long as *var* (*A|g*) */var* (*A|g* = 0) *≠* exp (*-ϖ*_*g*_ (*g*)).

A consistent estimator of *β*_*a*_ is therefore obtained as in the previous Section, by substituting in consistent estimators of unknown parameters and sample averages for expectations. To ground ideas, suppose that *-ϖ*_*g*_ (*g*) = *-ϖ*_*g*_*g* for vector *-ϖ*_*g*_, then a consistent estimator *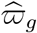* of *-ϖ*_*g*_ is given by the solution to the estimating equation:

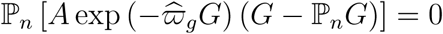

Note that if *A* is a rare binary exposure then *var* (*A|g*) */var* (*A|g* = 0) *≈* exp (*-ϖ*_*g*_ (*g*)) for all *g,* therefore violating the identification condition. In such instance, we recommend using the additive model described in the previous Section. For count data, the result rules out using a Poisson model for exposure, however other models that accommodate over-dispersion such as the negative binomial distribution may be used. Finally, it is straightforward to verify that the Lemma continues to hold if assumption 2 is dropped to allow for unmeasured confounding of the effects of *G* provided that the conditional covariance between the residual (*U*_*a*_*/E*(*U*_*a*_*|G*) *-* 1) and *U*_*y*_ given *G* does not depend on *G.* Note that in this latter case *E* (*A|G* = *g*) = exp (*-ϖ*_*g*_ (*g*)) = exp (*α*_*g*_ (*g*)) *E* (*U*_*a*_*|G* = *g*).

### 5.2 Incorporating Covariates

One may wish in an MR analysis to adjust for covariates, either to account for observed confounding of the exposure effect on the outcome, or to account for confounding of the effects of the genetic markers primarily by ancestry (known as population stratification) or simply to improve efficiency. In order to account for covariates *C*, we propose to solve:

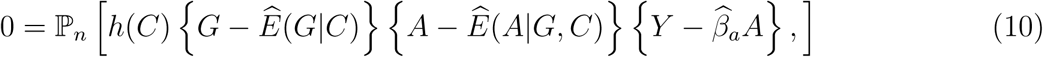

for user-specified choice of *h,* where Ê(*G|C*) and Ê(*A|G, C*) are consistent estimators of Ê(*A|G, C*) and Ê(*G|C*) obtained say by fitting appropriate generalized linear models. For example, as *G* is binary, one may specify logitPr(*G* = 1*|C*) = *ω*_0_ + *ω’ C* to obtain *Ê*(*G|C*) by standard likelihood estimation of a logistic regression, and likewise when *A* is binary, one may obtain *Ê*(*A|G, C*) by fitting a similar logistic regression, and when *A* is continuous, an analogous linear regression could be used instead. Identification results established in previous Sections continue to apply by further conditioning on *C* in Assumptions 1,2,2*,3,4, as well as on the left hand-side of equation (5). Note that effect modification can be incorporated upon conditioning on *C* in equation (1), by modeling the conditional causal effect of *A* on *Y* given *U* and *C* as a function of *C,* for example

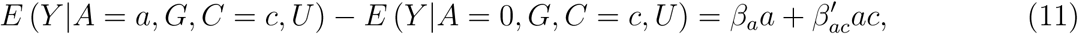

in which case *β*_*ac*_ captures effect modification by *C.* In contrast, effect modification by *C* in equation (2) can remain unrestricted, i.e.

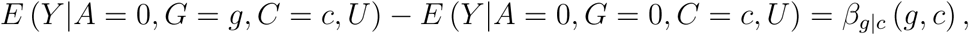

where *β*_*g|c*_ (0*, c*) = 0 for all *c* but is otherwise unrestricted. Estimation of (*β*_*a*_*, β*_*ac*_) requires modifying equation (10) as followed:

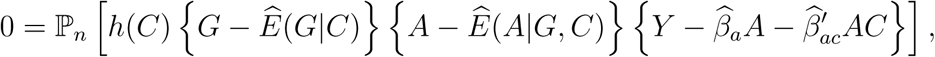

where *h*(*C*) is of the same dimension as (*A, C’*)*’,* e.g. *h*(*C*) = (1*, C’*)*’.*

### 5.3 Incorporating Multiple IVs

MR designs with multiple candidate genetic IVs may be used to strengthen identification and improve efficiency. Multiple candidate IVs can be incorporated by adopting a standard generalized method of moments approach. Specifically, suppose that *G* is a vector of genetic variants, then, assuming for simplicity that there is no effect modification of *A* by *C* in the outcome model, i.e.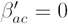, we propose to obtain 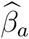 by solving:

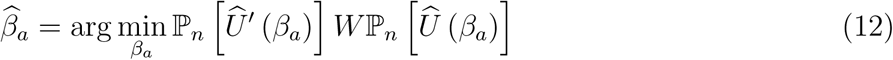

where

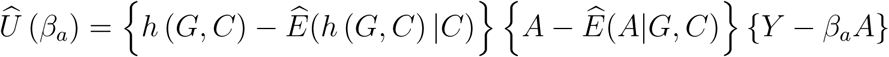

for a user-specified function *h* (*G, C*) of dimension *K* ≥ 1, and *W* is user-specified weight matrix. In practice, it may be convenient to set *h* (*G, C*) = *G* and *W* = *I*_*KxK*_ the *K* dimensional identity matrix. Let *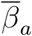* denote the corresponding estimator. A more efficient estimator *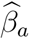* can then be obtained by solving (12) with weight *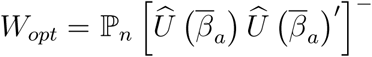* where *T*^−^ denotes the generalized inverse of matrix *T*. Identification of GMM is guaranteed (at least locally) provided that the second derivative wrt *β*_*a*_ of the GMM objective function 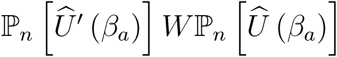 is nonsingular at the truth, which is a generalization of condition (5). The asymptotic distribution of *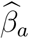* which solves (12) is described in Appendix A3.

### 5.4 Multiplicative causal effects

In this Section, we consider making inferences about the multiplicative causal effect of exposure *A,* under the model

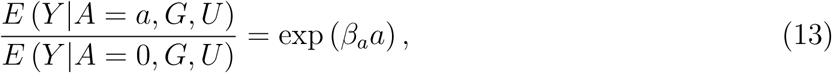

where for simplicity, we assume no baseline covariates, binary *A* and scalar *G*. Therefore, If *Y* is binary, *β*_*a*_*a* encodes the conditional log risk ratio log *{*Pr (*Y* = 1*|A* = *a, G, U*) */* Pr (*Y* = 1*|A* = 0*, G, U*)*}* which is assumed to be independent of *U* and *G,*i.e. there is no multiplicative interaction between *A* and (*G, U*). In order to state our identification result with an invalid IV, consider the following assumption.

**Assumption 5.** Equations (2), (3), and (13) hold.

#### Lemma 5

*Suppose that Assumptions 1,2*,5 hold, then β*_*a*_ *is the unique solution to equation:*

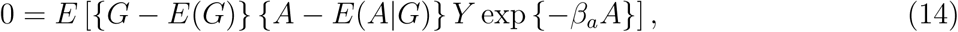

*provided that condition* (5) *holds.*

**Proof.** The results follows upon noting that *E* [*Y* exp *{-β*_*a*_*A} |A, G, U*] = *E* [*Y |A* = 0*, G, U*]. The proof then proceeds as in Lemma 1.

According to the Lemma, a consistent estimator of *β*_*a*_ can be obtained by solving an empirical version of equation (14) in a similar manner as in previous Sections. The unbiasedness property given by equation (14) continues to hold for continuous *A* under the conditions given in the Lemma, and generalizations to allow for covariates and multiple IVs can easily be deduced from previous Sections.

Interestingly, equation (14) continues to hold under case-control sampling wrt the outcome *Y*, however note that *E*(*G*) and *E*(*A|G*) must be evaluated wrt the underlying distribution for the target population which will in general not match the corresponding distributions in the casecontrol sample. To use the result in practice, one would either need to obtain these quantities from an external source or one could alternatively approximate them with the corresponding data distribution in the controls (i.e. units with *Y* = 0) provided the outcome is sufficiently rare. In the event sampling fractions for cases and controls are available, one could in principle implement inverse-probability of sampling weights to consistently estimate *E*(*G*) and *E*(*A|G*). Unbiasedness under case-control sampling follows from noting that *f* (*A, G, U|Y* = 1) ∝ Pr(*Y* = 1*|A, G, U*)*f* (*A, G, U*), and therefore

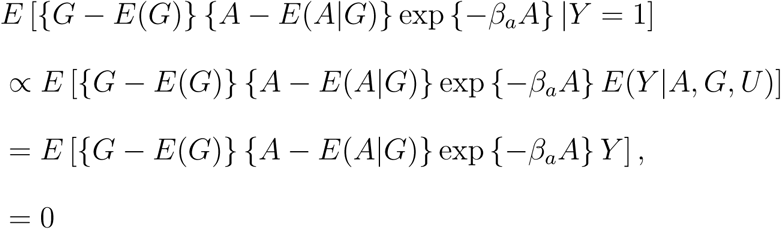

where the last equality follows from Lemma 3.

### 5.5 More efficient MR GENIUS

Similar to standard g-estimation, MR GENIUS can be made more efficient by incorporating information about the association between *G* and *Y.* This can be achieved by the following steps:

1. Obtain the MR GENIUS estimator 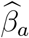 either on the additive or multiplicative scale.
2. Define a treatment-free outcome 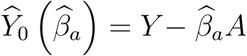 under (1) and 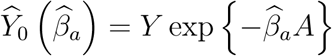 under (13).
3. Regress 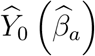 on *G* using a generalized linear model with appropriate link function, and define 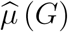 a person’s corresponding fitted (predicted) value.
4. Define 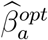 as the solution to

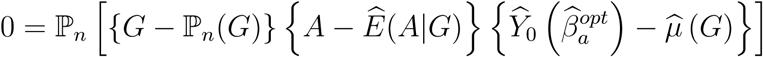

with 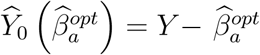 under (1) and 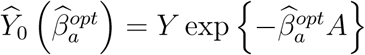 under (13).

If all regression models are correctly specified (including the glm for *E*(*Y*_0_ (*β*_*a*_) *|G*) required in Step 3 of the above procedure), a standard argument of semiparametric theory implies that the asymptotic variance of 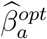 is guaranteed to be no larger than that of 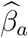 (Robins, 1997). Interestingly, MR GENIUS and its more efficient version coincide (up to asymptotic equivalence) whenever nonparametric methods are used to estimate all nuisance parameters, i.e. to estimate *E*(*G*)*, E*(*A|G*) and *µ* (*G*) = *E*(*Y - β*_*a*_*A|G*). For instance, in the case of binary *G,* such that regression models *E*(*A|G*) and *µ* (*G*) = *E*(*Y - β*_*a*_*A|G*) are saturated, the two estimators are exactly equal and yield identical inferences. Both approaches also coincide if all IVs are valid, however the above modification will tend to be more efficient with increasing number of invalid IVs. Note that *µ*(*G*) does not necessarily have a causal interpretation as the effect of *G* on *Y* may be confounded by *U*. Also note that misspecification of a model for *µ*(*G*) does not affect consistency and asymptotic normality of the MR GENIUS estimator of *β*_*a*_ provided that as we have assumed throughout, the model for *E*(*A|G*) is correct.

In the case of multiplicative outcome model, it is straightforward to extend the robustness properties of the efficient MR GENIUS estimator described above under an assumption of no multiplicative interaction (rather than no additive interaction) between *G* and *U.* This would simply entail replacing 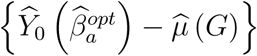 in step 4 with 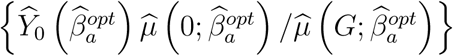 where 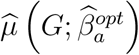 is the regression of 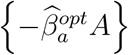 on *G* under an appropriate GLM and solving the estimating equation in Step 4 for 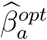. One can show using the same method of proof used throughout, that the resulting estimator is consistent for the causal effect of interest under violation of both assumptions 2 and 3, under an assumption analogous to Assumption 2*. Note however that 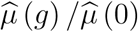 would now need to be consistent for *E*(*Y |A* = 0*, G* = *g*)*/E*(*Y |A* = 0*, G* = 0). It is likewise possible to modify the above procedure to accommodate a multiplicative exposure model by substituting in 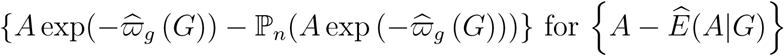 in Step 4.

### 5.6 Odds ratio exposure model

In this Section, we briefly consider how MR GENUIS might be applied in a setting where assumption 2 is replaced by the following weaker conditional independence assumption:

**Assumption 2**^†^. IV conditional independence: *G⊥ ⊥ U|A*;

A key implication of this assumption is that the causal effect of *G* on *Y,* is now identified conditional on *A,* because the assumption implies no unmeasured confounding of the effects of *G* on *Y.* Note however that *G* and *U* are not marginally independent. Suppose also that instead of assumption 4.b, one wishes to encode the IV-exposure association on the odds ratio scale, under the following homogeneity assumption:

**(4b**^†^) There is no odds ratio *G - U* interaction in model for *E* (*A|G, U*)

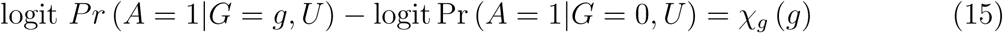

for an unknown function *χ*_*g*_ (*·*) that satisfies χ_*g*_ (0) = 0.

We then have the following identification result for the multiplicative causal effect *β*_*a*_ of model (13).

#### Lemma 5

*Under Assumptions 1.2†, 4.b† and equations* (2) *and* (13) *, we have that β*_*a*_ = *θ, where θ is the unique solution to equation:*

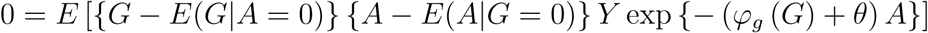

*where*

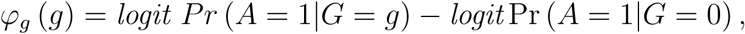

*provided that γ*_*ag*_(*g*) *≠* 0 *for some value of g, with:*

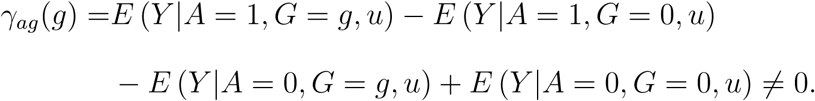

Assumption 2^†^ in fact implies that *ϕ*_*g*_(.) = *χ*_*g*_(.) (Ma et al 2006). The Lemma establishes that under Assumptions 1, 2^†^, 4.b^†^ and equations (2) and (13), the multiplicative causal effect of *A* is identified, provided that *γ*_*ag*_(*g*) *≠* 0. In the proof of the Lemma given in Appendix A1, we establish that under our assumptions *γ*_*ag*_(*g*) = (exp (*β*_*a*_) *-* 1) *β*_*g*_ (*g*), and therefore the causal effect is not identified by the Lemma if all IVs satisfy the exclusion restriction assumption, such that *β*_*g*_ (*g*) = 0 for all *g.* Note that the latter assumption is empirically testable because the direct effect of *G* on *Y* is unconfounded. If *β*_*g*_(*g*) =*/* 0 for some *g*, a valid test for the causal null hypothesis can be performed by testing whether the estimating equation given in the Lemma holds at *θ* = 0. An estimator of *β*_*a*_ based on the Lemma is easily deduced from previous Sections.

### 5.7 MR GENUIS for censored failure time under a multiplicative survival model

Censored time-to-event endpoints are common in MR studies and IV methods to address such data are increasingly of interest; recent contributions to this literature include Nie et al (2011), Tchetgen Tchetgen et al (2015), Li et al (2015) and Martinussen et al (2017). While these methods have been shown to produce a consistent causal effect estimator encoded either on the scale of survival probabilities, or as a hazards ratio or hazards difference, leveraging a valid IV which satisfies assumptions (1)-(3), they are not robust to violation of any of these assumptions. In this Section, we briefly extend MR GENIUS to survival analysis under an additive hazards model. Thus, suppose now that *Y* is a time-to-event outcome which satisfies the following additive hazards model

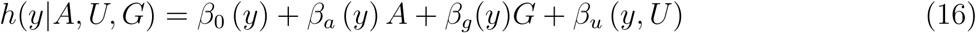

where *h*(*y|A, U, G*) is the hazard function of *Y* evaluated at *y*, conditional on *A, U* and *G*, and the functions (*β*_0_ (*·*) *, β*_*a*_ (*·*) *, β*_*g*_(*·*)*, β*_*u*_ (*·, ·*)) are unrestricted. The model states that conditional on *U*, the effect of *A* on *Y* encoded on the additive hazards scale is linear in *A* for each *y*, although, the effect size *β*_*a*_ (*y*) may vary with *y*. The model is quite flexible in the unobserved confounder association with the outcome *β*_*u*_ (*·, ·*), which is allowed to remain unrestricted at each time point*y* and across time points. This is the model considered by Tchetgen Tchetgen et al (2015) who further assumed that *β*_*g*_(*y*) = 0 for all *y* by the exclusion restriction assumption 3. Here we do not make this assumption. As usually the case in survival analysis, *Y* is subject to right-censoring due to drop-out, and therefore instead of observing *Y* for all subjects, one observes *Y \** = min(*Y, X*) and Δ = *I*(min(*Y, X*) = *Y*), where *X* is an independent censoring time (i.e. independent of *Y, A, G, U*). Let *R*(*y*) = *I*(*Y * ≥ y*) denote the at-risk process and *N* (*y*) = *I*(*Y * ≤ y,* Δ = 1) the counting process associated with failure time. As discussed in Martinussen et al (2017), the additive hazards model (16) is particularly attractive because it implies a multiplicative survival model for the joint causal effect of *A* and *G* on *Y*:

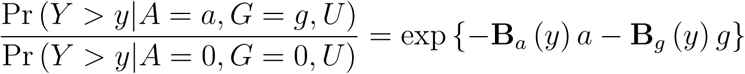

where 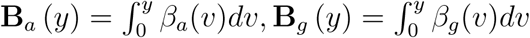. Our objective is therefore to identify and estimate **B**_*a*_ (*y*). We have the following result which extends the result of Martinussen et al (2017) in order to accommodate possible violation of the exclusion restriction assumption:

#### Lemma 6

*Under assumptions 1,2,4.b and equation* (16) *, we have that for each y*

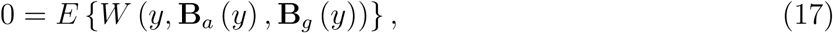

*where*

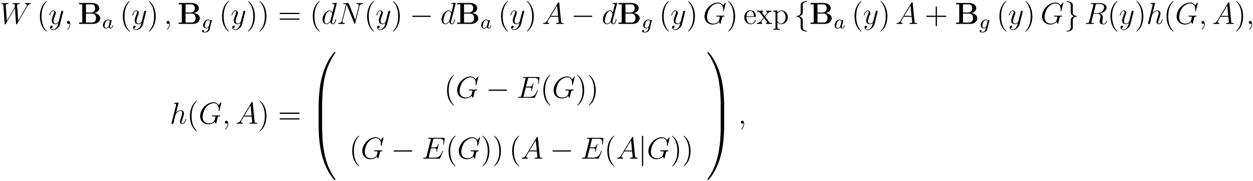

**Proof.** We note that by assumption

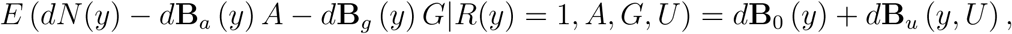

and

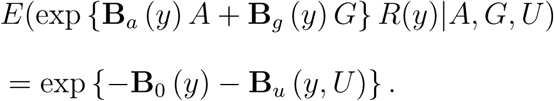

Therefore

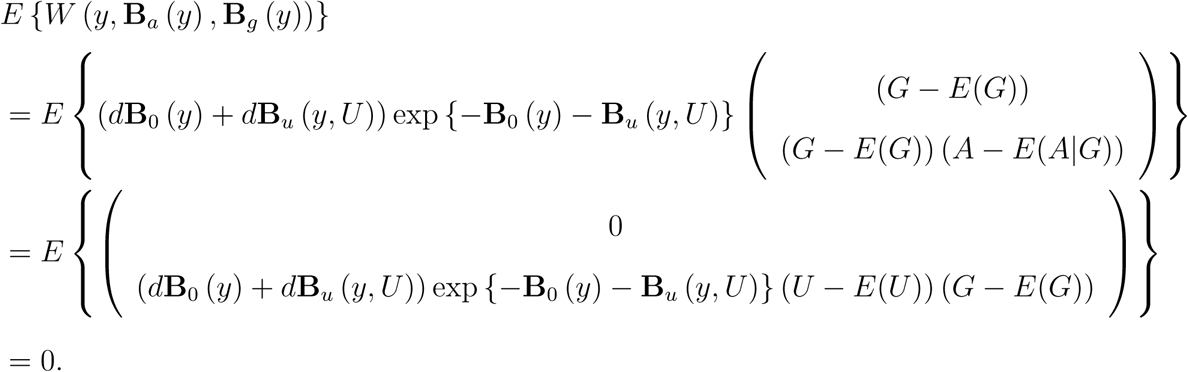

As in Martinussen et al (2017), the unbiasedness of equation *W* (*y,* **B**_*a*_ (*y*), **B**_*g*_ (*y*)) suggests a way of estimating the increments (*d***B**_*a*_ (*y*) *, d***B**_*g*_ (*y*)) by solving an empirical version of equation (17) for each *y* with population expectations replaced by sample analogs, giving the following recursive estimator

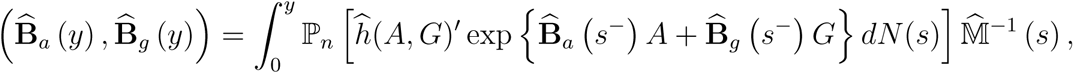

where 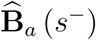 is the value of 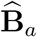 right prior to *s*, and likewise for 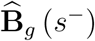, and

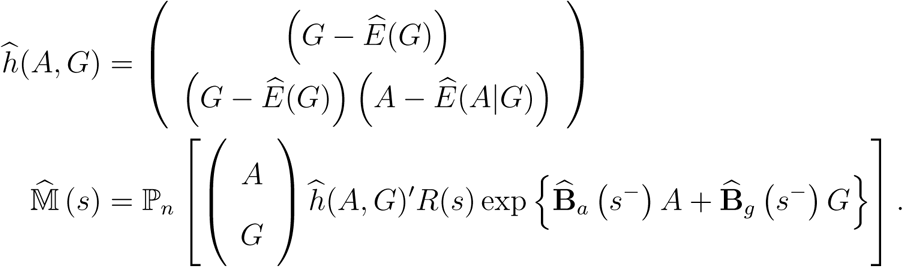

Because of its recursive structure, this estimator can be solved forward in time starting with (*d***B**_*a*_ (0) *, d***B**_*g*_ (0)) = (0, 0). The resulting estimator is a counting process integral, therefore only changing values at observed event time. The estimator is only defined provided M (*y*) is invertible at each such jump time, which is essentially a necessary condition for identification. The large sample behavior of the resulting estimator follows from results derived in Martinussen et al (2017) and is therefore omitted. Note that the result relies on assumption 2 therefore ruling out confounding of the effect of the IV on the outcome.

## 6 Simulation Study

### 6.1 Single IV

We investigate the finite-sample properties of MR GENIUS proposed above and compare them with existing estimators under a variety of settings. For a single binary IV *G*, we generate independent and identically distributed (*G*_*i*_*, U*_*i*_*, A*_*i*_*, Y*_*i*_), *i* = 1, 2*, …, n* as follows:

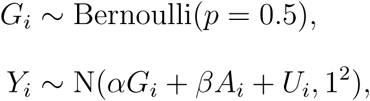

where for binary exposure *A*,

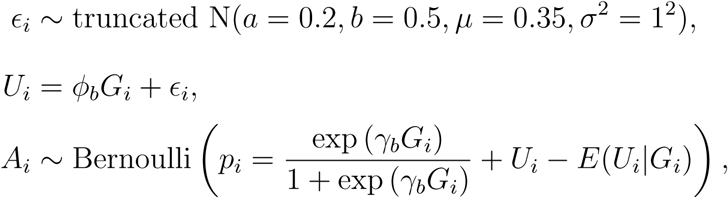

where *∊*_*i*_ is appropriately bounded to ensure that *p* falls in the unit interval, and for continuous *A*,

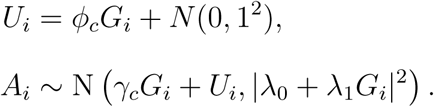

The data generating mechanism satisfies assumptions 2* and 4. We set *γ*_*b*_ = *-*0.5 or *-*1 (binary *A*), and *γ*_*c*_ = *-*1, *λ*_0_ = 1, *λ*_1_ = 1 or 5 (continuous *A*) which satisfy both Assumption 1 and condition (5). Assumptions 2 and 3 are violated when we set *φ*_*b*_ = *-*0.2, *φ*_*c*_ = *-*2 and *α* = *-*0.5 respectively. The causal parameter is set equal to *β* = 0.5 throughout this simulation. The IV strength is tuned by varying the values of *γ*_*b*_ and *λ*_1_, for binary and continuous *A* respectively.

MR GENIUS is implemented using (6), with *Ê*(*A|G*) estimated with linear or logistic regression when *A* is continuous or binary, respectively. In this single-IV setting, we also implement the twostage least squares (TSLS) estimator, which is the most common approach used in practice. The simulation results based on 1000 replicates at sample sizes *n* = 500 and *n* = 1000 are summarized in Tables 1 and 2, for continuous and binary exposure respectively. When Assumptions 2 and 3 both hold, TSLS and MR GENIUS have small bias regardless of sample size. Coverage of the Wald-type 95% confidence interval (CI) for the causal parameter is also close to nominal level. Efficiency of the estimators increases with IV strength. When the IV is invalid, TSLS is biased and its 95% CI undercovers, while in accordance with theory MR GENIUS continues to have small bias and correct coverage.

**Table 1:** Monte Carlo results of MR GENIUS and TSLS estimation of *β*_0_ = 0.5 with continuous exposure and single IV at two different strengths (*λ*_1_ = 1, 5). The first and second rows’ results for each estimator correspond to sample sizes *n* = 500, 1000 respectively.

| | | $\vert \lambda_1\vert = 1$ | | | $\vert \lambda_1\vert = 5$ | | |
| --- | --- | --- | --- | --- | --- | --- | --- |
|  |  | TTT<sup>†</sup> | TTF | TFF | TTT | TTF | TFF |
| Median absolute value of bias |  |  |  |  |  |  |  |
| MR GENIUS |  | 0.00 | 0.00 | 0.00 | 0.00 | 0.00 | 0.00 |
|  |  | 0.00 | 0.00 | 0.00 | 0.00 | 0.00 | 0.00 |
| TSLS |  | 0.00 | 0.50 | 0.83 | 0.00 | 0.51 | 0.84 |
|  |  | 0.00 | 0.50 | 0.83 | 0.00 | 0.50 | 0.83 |
| Median estimated SD |  |  |  |  |  |  |  |
| MR GENIUS |  | 0.08 | 0.08 | 0.08 | 0.02 | 0.02 | 0.02 |
|  |  | 0.06 | 0.06 | 0.06 | 0.01 | 0.01 | 0.01 |
| TSLS |  | 0.13 | 0.12 | 0.05 | 0.14 | 0.22 | 0.11 |
|  |  | 0.09 | 0.09 | 0.03 | 0.09 | 0.15 | 0.08 |
| Monte Carlo SD<sup>‡</sup> |  |  |  |  |  |  |  |
| MR GENIUS |  | 0.08 | 0.08 | 0.08 | 0.02 | 0.02 | 0.02 |
|  |  | 0.06 | 0.06 | 0.06 | 0.01 | 0.01 | 0.01 |
| TSLS |  | 0.12 | 0.13 | 0.05 | 0.13 | 0.25 | 0.11 |
|  |  | 0.09 | 0.09 | 0.04 | 0.09 | 0.15 | 0.08 |
| 95% Wald-type CI coverage |  |  |  |  |  |  |  |
| MR GENIUS |  | 95.3 | 95.3 | 95.3 | 94.8 | 94.8 | 94.8 |
|  |  | 94.8 | 94.8 | 94.8 | 95.2 | 95.2 | 95.2 |
| TSLS |  | 95.3 | 2.7 | 0.0 | 99.0 | 36.9 | 0.0 |
|  |  | 95.9 | 0.1 | 0.0 | 97.7 | 5.8 | 0.0 |

<sup>†</sup>: TTT: IV assumptions (1), (2) and (3) hold; TTF: IV assumption (3) (exclusion restriction) does not hold; TFF: both IV assumptions (2) and (3) (IV independence) do not hold.
<sup>‡</sup>: Robust normal-consistent estimate obtained from dividing the interquartile range of causal effect estimates by 1.349.

**Table 2:** Monte Carlo results of MR GENIUS and TSLS estimation of *β*_0_ = 0.5 with binary exposure and single IV at two different strengths (*γ*_*b*_ = *−*0.5*, −*1). The first and second rows’ results for each estimator correspond to sample sizes *n* = 500, 1000 respectively.

| | | $\vert \gamma_b\vert = 0.5$ | | | $\vert \gamma_b\vert = 1$ | | |
| --- | --- | --- | --- | --- | --- | --- | --- |
|  |  | TTT<sup>†</sup> | TTF | TFF | TTT | TTF | TFF |
| Median absolute value of bias |  |  |  |  |  |  |  |
|  | MR GENIUS | 0.01 | 0.01 | 0.01 | 0.03 | 0.03 | 0.01 |
|  |  | 0.01 | 0.01 | 0.00 | 0.01 | 0.01 | 0.00 |
|  | TSLS | 0.00 | 4.05 | 2.16 | 0.00 | 2.17 | 1.61 |
|  |  | 0.02 | 4.07 | 2.18 | 0.01 | 2.18 | 1.61 |
| Median estimated SD |  |  |  |  |  |  |  |
|  | MR GENIUS | 0.54 | 0.54 | 0.33 | 0.36 | 0.36 | 0.34 |
|  |  | 0.37 | 0.37 | 0.23 | 0.25 | 0.25 | 0.24 |
|  | TSLS | 0.77 | 1.39 | 0.38 | 0.40 | 0.53 | 0.25 |
|  |  | 0.53 | 0.97 | 0.27 | 0.28 | 0.37 | 0.18 |
| Monte Carlo SD<sup>‡</sup> |  |  |  |  |  |  |  |
|  | MR GENIUS | 0.54 | 0.54 | 0.32 | 0.35 | 0.35 | 0.34 |
|  |  | 0.39 | 0.39 | 0.22 | 0.25 | 0.25 | 0.24 |
|  | TSLS | 0.79 | 1.37 | 0.37 | 0.41 | 0.52 | 0.27 |
|  |  | 0.55 | 1.06 | 0.28 | 0.29 | 0.37 | 0.18 |
| 95% Wald-type CI coverage |  |  |  |  |  |  |  |
|  | MR GENIUS | 98.7 | 98.7 | 96.0 | 95.2 | 95.2 | 95.8 |
|  |  | 96.9 | 96.9 | 96.4 | 95.9 | 95.9 | 94.9 |
|  | TSLS | 98.6 | 9.7 | 0.0 | 95.2 | 0.0 | 0.0 |
|  |  | 96.4 | 0.3 | 0.0 | 94.6 | 0.0 | 0.0 |

<sup>†</sup>: TTT: IV assumptions (1), (2) and (3) hold; TTF: IV assumption (3) (exclusion restriction) does not hold; TFF: both IV assumptions (2) and (3) (IV independence) do not hold.
<sup>‡</sup>: Robust normal-consistent estimate obtained from dividing the interquartile range of causal effect estimates by 1.349.

#### Multiple IVs

Here we generate i.i.d. *L*_*i*_ = (*G*_*i*_*, U*_*i*_*, A*_*i*_*, Y*_*i*_), *i* = 1, 2*, …, n*, with *p*_*G*_ = 10 IVs from:

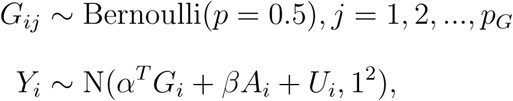

where *G*_*i*_ = (*G*_*i*1_*, G*_*i*2_*, …, G*_*ipG*_)^*T*^. For binary exposure,

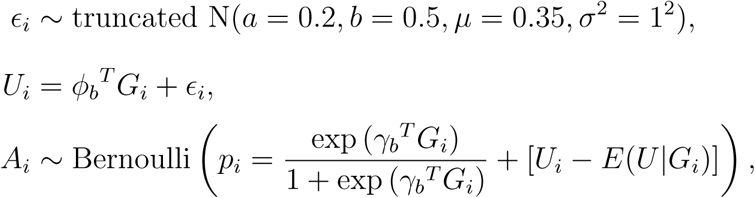

where *∊*_*i*_ is appropriately bounded to ensure that *p*_*i*_ falls in the unit interval, and for continuous exposure,

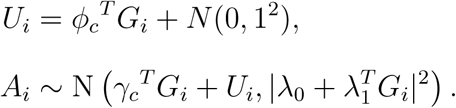

For binary exposure, IV strength is set to *-*0.15 for each entry of *γ*_*b*_, while in the continuous exposure case each entry of *γ*_*c*_ and *λ*_1_ is set identically to *-*2 and 0.5 respectively. We first generate an ideal scenario in which all 10 IVs are valid and satisfy Assumptions 1-3, next we consider scenarios where the first three, six or all of the IVs are invalid. With three invalid IVs, *α*^*T*^ = *-*0.5 *·* (1, 1, 1, 0*, …,* 0) and 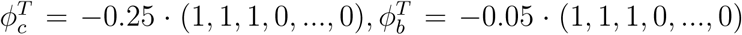 when Assumption 3 or 2 is violated, respectively; with six invalid IVs, *α*^*T*^ = *-*0.25(1, 1, 2, 2, 4, 4, 0*, …,* 0) and 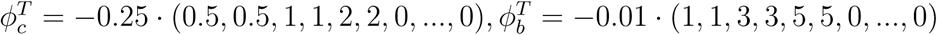 accordingly. When all IVs are invalid, *α*^*T*^ = *-*0.5 *·* (1, 1*, …,* 1) and 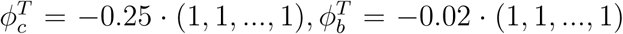. The setting with three invalid IVs investigates the condition in which fewer than 50% of the IVs are invalid (Kang et al, 2016; Windmeijer et al, 2016); in the setting with six invalid IVs this condition is violated, but the set of valid IVs form the largest group according to the plurality rule (Guo et al, 2017).

MR GENIUS is implemented as the solution to (12) with optimal weight; the more efficient version of MR GENIUS as described in section 5.4 is also implemented. MR-Egger regression estimation, TSLS (which assumes all IVs are valid) and sisVIVE are implemented using the R packages MendelianRandomization, AER and sisVIVE (Yavorska and Burgess, 2017; Kleiber and Zeileis, 2008; Kang, 2017) respectively, under default settings. The adaptive Lasso and TSHT estimation methods are implemented as described in Windmeijer et al (2016) and Guo et al (2017) respectively. We also implement post-adaptive Lasso which uses adaptive Lasso for the purpose of selecting valid IVs but not in the process of estimating the causal effect. We also implement the oracle TSLS which assumes the set of valid IVs to be known a priori.

Simulation results based on 1000 replications for sample sizes of *n* = 500, 2000 and 10, 000 with continuous exposure are presented in Tables 3 and 4. When there are zero or three invalid IVs (majority rule holds), the sisVIVE, adaptive, post-adaptive Lasso and TSHT estimators exhibit small bias which becomes negligible at sample size of *n* = 10, 000. Empirical coverage of CIs is close to nominal level once *n* ≥ 2000 for adaptive/post-adaptive Lasso and TSHT. Adaptive Lasso and TSHT on average correctly identifies invalid IVs, while sisVIVE on average selects four IVs as invalid when there are three in truth (see Table 7 for results on IV selection). The naive TSLS estimator performs well in terms of bias and coverage only when all IVs are valid; as expected, it is biased and its 95% CI severely undercovers in all other settings with at least one invalid IV. Post-adaptive Lasso is generally less biased in finite sample than adaptive Lasso. Post-adaptive Lasso and oracle TSLS perform similarly in terms of bias and efficiency once *n* ≥ 2000 (when the majority rule holds), in agreement with theory since they are asymptotically equivalent under these settings. MR GENIUS also has small bias at all sample sizes and its bias becomes negligible at *n* = 10000, with adequate 95% CI empirical coverage at *n* ≥ 500. MR GENIUS is generally less efficient than the other estimators when the majority rule holds, except for MR-Egger. MR-Egger exhibits some bias, but its 95% CI coverage is adequate when there are no invalid IVs, with slight undercoverage when there are three invalid IVs. Since MR-Egger assumes a two-sample analysis whereby association coefficients relating IVs and exposure/outcome are uncorrelated, the observed bias may be a reflection of the single sample simulation setting.

**Table 3:**
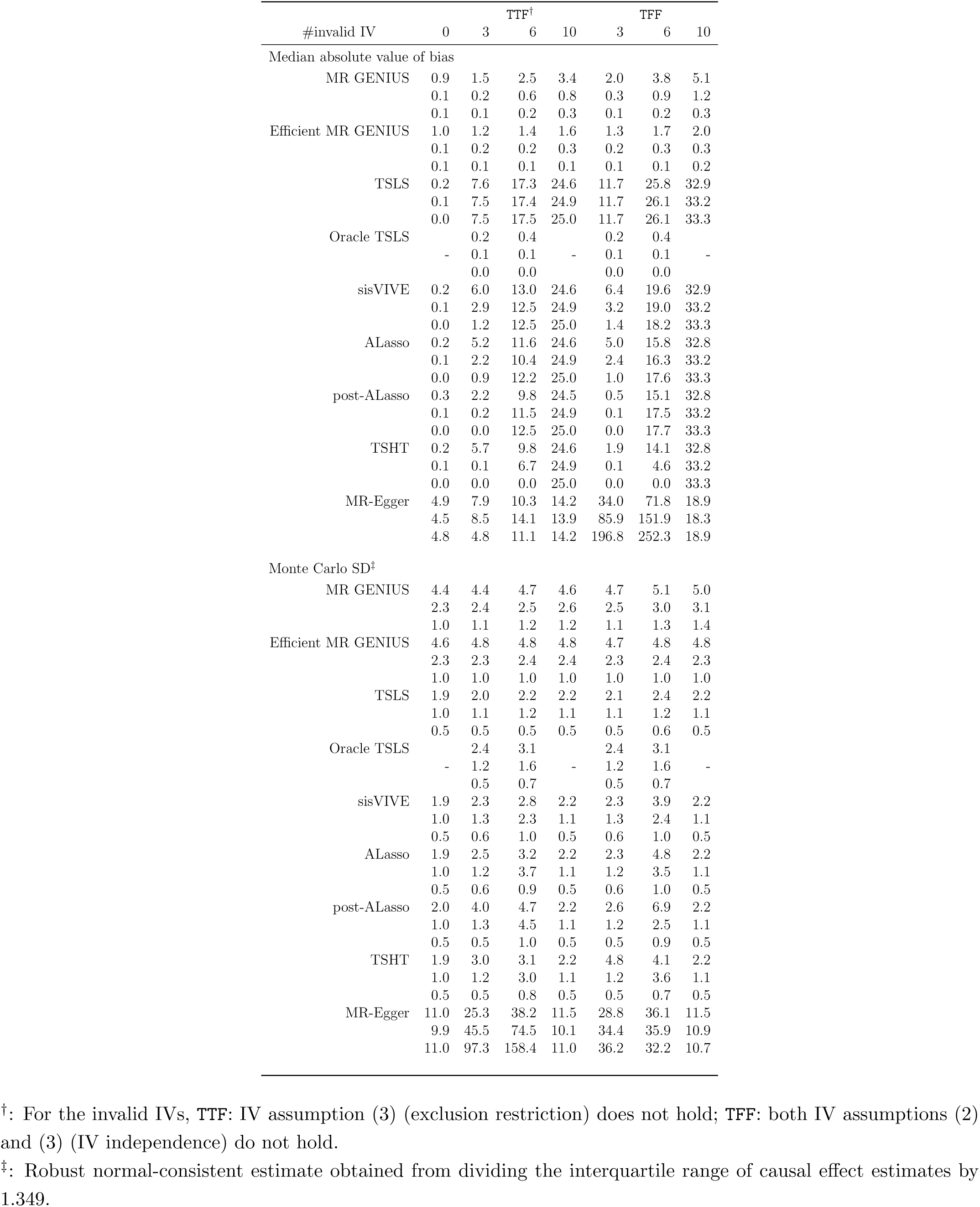
Median absolute value of bias and Monte Carlo standard error in estimation of *β*_0_ = 0.5 with continuous exposure and *p*_*G*_ = 10 IVs. All entries are original values multiplied by 100. The three rows of results for each estimator correspond to sample sizes of *n* = 500, *n* = 2000 and *n* = 10, 000 respectively.

**Table 4:**
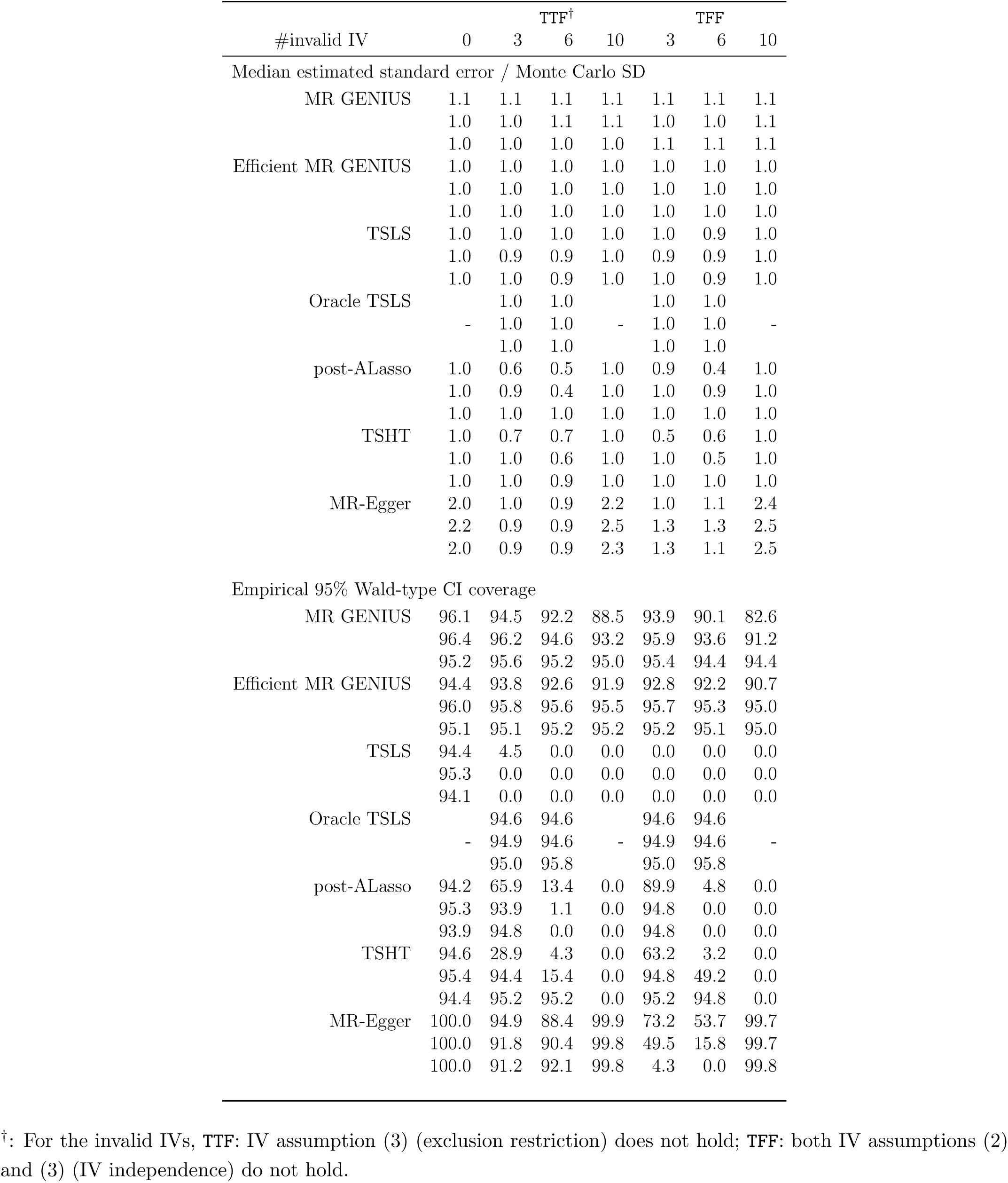
Ratio of estimated to Monte Carlo standard error and empirical 95% Wald-type CI coverage in estimation of *β*_0_ = 0.5 with continuous exposure and *p*_*G*_ = 10 IVs. The three rows of results for each estimator correspond to sample sizes of *n* = 500, *n* = 2000 and *n* = 10, 000 respectively. Only point estimation is implemented for sisVIVE and adaptive Lasso, hence their results are not available.

When six IVs are invalid and the majority rule is violated, sisVIVE and adaptive/post-adaptive Lasso are significantly biased, with no improvement as sample size increases. There is also increasing undercoverage of 95% CI as sample size increases for post-adaptive Lasso. On average, sisVIVE and adaptive Lasso select eight IVs as invalid when only six are actually invalid, and fails to select any IV as invalid when all are. TSHT is also biased and its 95% CI undercovers when all IVs are invalid (with none of the IVs selected as invalid on average in this case); however when six IVs are invalid, the plurality rule holds and its bias diminishes at *n* = 10, 000. Adequate 95% CI coverage is also achieved at *n* = 10, 000, with the right number of IVs selected as invalid on average. The efficiency of estimators generally decreases with increasing number of invalid IVs, except when all the IVs are invalid. The bias of MR GENIUS improves with increasing sample size when six or all IVs are invalid; adequate 95% CI coverage is achieved at *n* ≥ 2000 with six invalid IVs, andat *n* = 10, 000 with all IVs invalid. Efficiency comparisons MR GENIUS is again generally less efficient than TSHT when six IVs are invalid, however it outperforms MR-Egger even when the InSIDE assumption holds (Bowden et al, 2015). However, MR-Egger is generally more biased, with severe 95% CI undercoverage, when the invalid IVs violate both Assumptions 2 and 3, which corresponds to a violation of the InSIDE assumption. The efficient MR GENIUS is generally less biased and more efficient compared to MR GENIUS, especially when more IVs are invalid. The estimated asymptotic relative efficiency of efficient MR GENIUS to MR GENIUS is approximately 0.5 with 10 invalid IVs (which violate both assumptions 2 and 3); the 95% CIs based on efficient MR GENIUS also attain correct coverage across all the scenarios at *n* ≥ 2000.

Simulation results with a binary exposure are summarized in Tables 5 and 6; the conclusions are mostly qualitatively similar to those in the continuous exposure setting. However, when there are six invalid IVs, TSHT is biased and its 95% CI undercovers, with no improvement as sample size increases. While the exposure is generated under a logit model (upon marginalizing over *U*), TSHT assumes a linear model which is misspecified in this simulation study. In addition, because the exposure is binary, most if not all IVs are weakly associated with *A* on the additive scale. Weak IVs may not be selected as valid IVs in the first thresholding step of TSHT (the number of IVs selected as relevant is nine on average at *n* = 10, 000); even if they are included, their inclusion may lead to incorrect inference in the subsequent estimation step (the number of IVs selected as relevant but invalid is close to three on average, when six are valid at *n* = 10, 000). MR-Egger appears to be less biased compared to the continuous exposure setting; this may be due to smaller values of *φ*_*b*_ used for binary exposure setting, so that violation of the InSIDE assumption is less severe.

**Table 5:** Median absolute value of bias and Monte Carlo standard error in estimation of *β*_0_ = 0.5 with binary exposure and *p*_*G*_ = 10 IVs. All entries are original values multiplied by 100. The three rows of results for each estimator correspond to sample sizes of *n* = 500, *n* = 2000 and *n* = 10, 000 respectively.

| #invalid IV | 0 | TTF<sup>†</sup> |  |  | TFF |  |  |
| --- | --- | --- | --- | --- | --- | --- | --- |
|  |  | 3 | 6 | 10 | 3 | 6 | 10 |
| Median absolute value of bias |  |  |  |  |  |  |  |
| MR GENIUS | 0.7 | 14.6 | 28.2 | 35.8 | 16.3 | 29.6 | 36.6 |
|  | 0.4 | 6.3 | 12.4 | 15.1 | 7.0 | 13.3 | 16.4 |
|  | 0.4 | 0.9 | 2.7 | 3.3 | 1.0 | 2.8 | 3.7 |
| Efficient MR GENIUS | 2.3 | 1.6 | 1.0 | 0.1 | 1.2 | 0.9 | 0.2 |
|  | 1.0 | 0.9 | 0.0 | 0.8 | 0.8 | 0.2 | 0.9 |
|  | 0.3 | 0.4 | 0.5 | 0.7 | 0.4 | 0.6 | 0.7 |
| TSLS | 1.0 | 177.9 | 433.4 | 616.8 | 197.0 | 455.3 | 642.0 |
|  | 0.1 | 328.3 | 788.9 | 1,112.9 | 361 | 828.2 | 1,156.9 |
|  | 0.6 | 431.6 | 1,006.4 | 1,436.1 | 474.1 | 1,057.7 | 1,493.4 |
| Oracle TSLS |  | 3.7 | 0.6 |  | 3.7 | 0.6 |  |
|  | - | 0.4 | 1.2 | - | 0.4 | 1.2 | - |
|  |  | 0.4 | 0.7 |  | 0.4 | 0.7 |  |
| sisVIVE | 1.0 | 82.9 | 302.1 | 616.2 | 81.9 | 311.0 | 642.0 |
|  | 0.2 | 75.5 | 568.6 | 1,111.1 | 74.5 | 590.2 | 1,156.7 |
|  | 0.6 | 46.9 | 716.3 | 1,435.6 | 46.8 | 747.8 | 1,492.7 |
| ALasso | 0.9 | 56.0 | 217.1 | 616.2 | 54.8 | 221.0 | 641.8 |
|  | 0.1 | 45.7 | 468.4 | 1,112.2 | 44.7 | 482.7 | 1,156.7 |
|  | 0.7 | 29.2 | 585.1 | 1,434.5 | 29.1 | 611.0 | 1,492.0 |
| post-ALasso | 0.8 | 13.1 | 180.0 | 615.8 | 11.8 | 181.7 | 641.3 |
|  | 1.0 | 0.8 | 395.0 | 1,111.1 | 0.4 | 408.8 | 1,156.0 |
|  | 0.7 | 0.2 | 558.7 | 1,432.7 | 0.2 | 579.8 | 1,490.7 |
| TSHT | 1.6 | 97.0 | 322.2 | 514.1 | 97.6 | 340.3 | 531.3 |
|  | 2.4 | 15.1 | 270.0 | 874.3 | 14.8 | 273.2 | 909.9 |
|  | 0.4 | 0.2 | 408.3 | 1,401.2 | 0.2 | 424.8 | 1,456.9 |
| MR-Egger | 8.8 | 165.1 | 339.3 | 447.4 | 182.5 | 356.8 | 463.8 |
|  | 2.4 | 133.6 | 333.8 | 171.7 | 148.0 | 353.4 | 175.6 |
|  | 0.4 | 10.8 | 45.2 | 17.9 | 15.1 | 47.7 | 17.7 |
| Monte Carlo SD<sup>‡</sup> |  |  |  |  |  |  |  |
| MR GENIUS | 69.3 | 76.1 | 91.4 | 88.5 | 76.9 | 93.4 | 89.7 |
|  | 48.2 | 54.5 | 65.9 | 69.1 | 55.3 | 66.6 | 71.0 |
|  | 26.3 | 28.7 | 35.0 | 35.9 | 29.5 | 35.5 | 36.4 |
| Efficient MR GENIUS | 67.9 | 68.4 | 67.8 | 67.4 | 68.4 | 67.7 | 67.1 |
|  | 48.4 | 48.2 | 48.2 | 48.7 | 48.2 | 48.3 | 48.8 |
|  | 26.1 | 26.1 | 26.3 | 26.1 | 26.1 | 26.3 | 26.1 |
| TSLS | 52.9 | 141.8 | 240.0 | 186.0 | 151.7 | 253.0 | 192.9 |
|  | 36.0 | 131.0 | 204.0 | 185.1 | 143.9 | 212.8 | 191.3 |
|  | 18.1 | 75.8 | 129.8 | 119.1 | 82.9 | 136.3 | 123.3 |
| Oracle TSLS |  | 65.0 | 89.0 |  | 65.0 | 89.0 |  |
|  | - | 46.8 | 60.7 | - | 46.8 | 60.7 | - |
|  |  | 22.3 | 29.2 |  | 22.3 | 29.2 |  |
| sisVIVE | 52.8 | 101.9 | 231.9 | 187.2 | 101.9 | 244.9 | 195.3 |
|  | 36.0 | 59.7 | 266.6 | 187.6 | 59.7 | 277.1 | 192.5 |
|  | 18.1 | 24.5 | 172.2 | 121.7 | 24.6 | 182.4 | 126.2 |
| ALasso | 52.8 | 70.6 | 210.1 | 186.1 | 65.3 | 219.7 | 193.5 |
|  | 36.0 | 38.9 | 261.4 | 184.5 | 38.1 | 285.4 | 191.6 |
|  | 18.0 | 22.0 | 170.0 | 121.4 | 21.9 | 186.4 | 125.9 |
| post-ALasso | 52.3 | 75.6 | 206.5 | 185.5 | 70.0 | 214.2 | 191.7 |
|  | 35.3 | 47.8 | 240.4 | 187.5 | 47.9 | 259.6 | 193.2 |
|  | 18.0 | 22.3 | 186.2 | 123.4 | 22.3 | 204.7 | 128.7 |
| TSHT | 64.2 | 221.6 | 391.0 | 193.2 | 239.5 | 414.7 | 201.7 |
|  | 48.4 | 69.6 | 463.9 | 125.2 | 68.5 | 498.4 | 128.8 |
|  | 18.1 | 22.5 | 222.5 | 115.8 | 22.5 | 236.4 | 119.8 |
| MR-Egger | 101.2 | 330.2 | 559.7 | 409.1 | 363.6 | 590.7 | 427.1 |
|  | 78.4 | 508.6 | 898.7 | 590.6 | 558.5 | 947.6 | 614.5 |
|  | 73.8 | 1,055.8 | 1,608.0 | 94.8 | 1,161.4 | 1,693.6 | 95.6 |

<sup>†</sup>: For the invalid IVs, TTF: IV assumption (3) (exclusion restriction) does not hold; TFF: both IV assumptions (2) and (3) (IV independence) do not hold.
<sup>‡</sup>: Robust normal-consistent estimate obtained from dividing the interquartile range of causal effect estimates by 1.349.

**Table 6:**
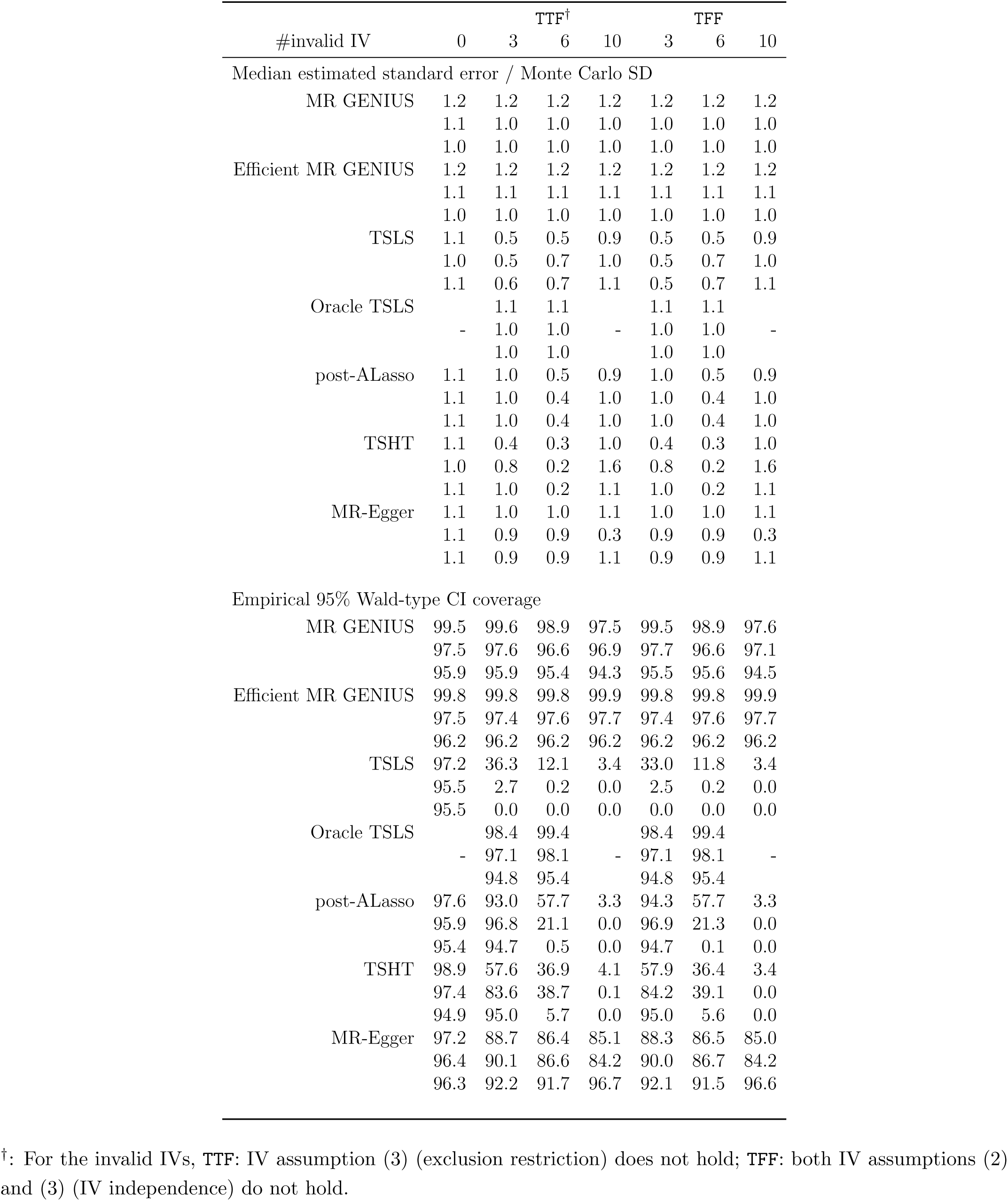
Ratio of estimated to Monte Carlo standard error and empirical 95% Wald-type CI coverage in estimation of *β*_0_ = 0.5 with binary exposure and *p*_*G*_ = 10 IVs. The three rows of results for each estimator correspond to sample sizes of *n* = 500, *n* = 2000 and *n* = 10, 000 respectively. Only point estimation is implemented for sisVIVE and adaptive Lasso, hence their results are not available.

**Table 7:**
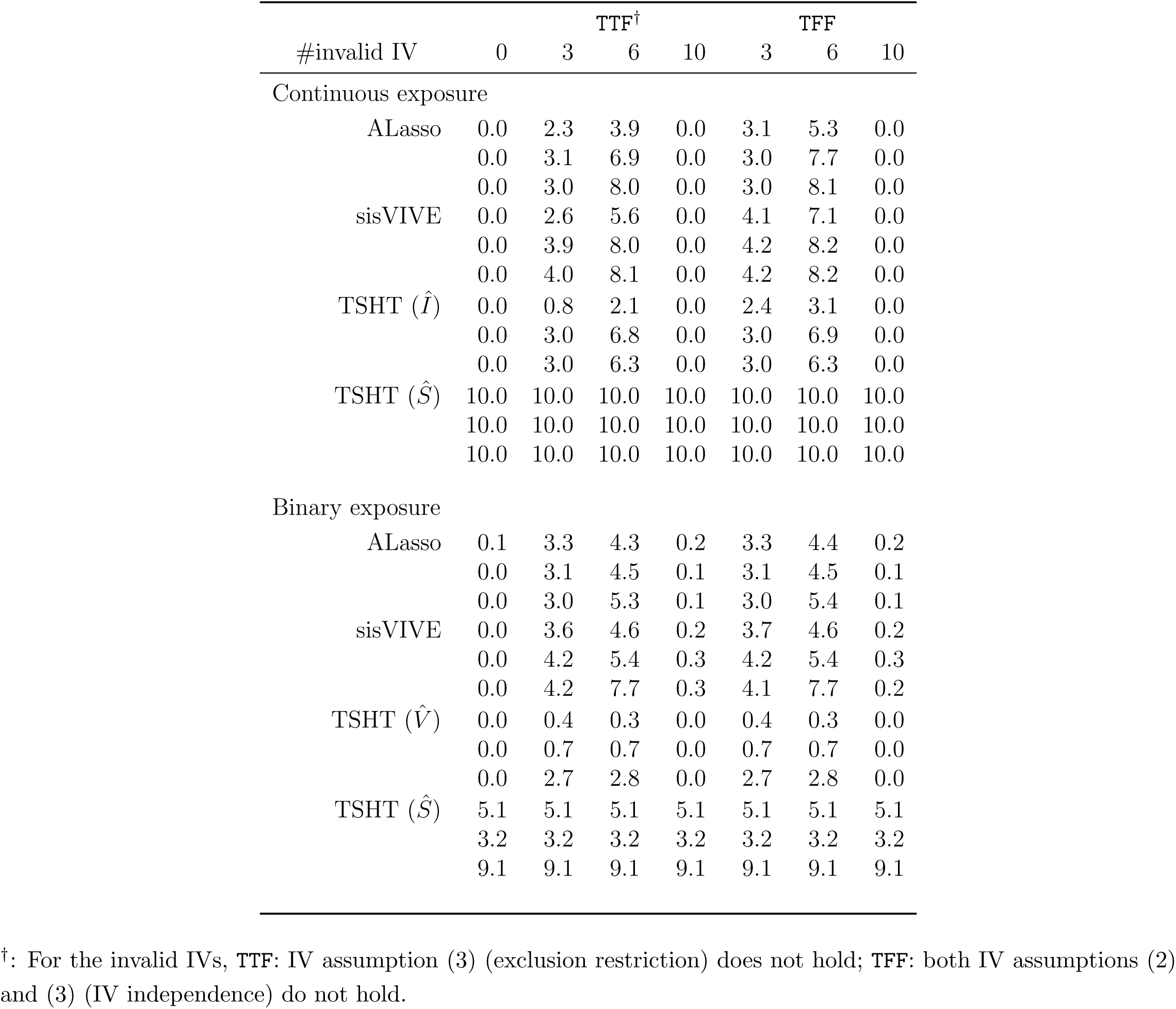
Average number of IVs selected as invalid by adaptive Lasso and sisVIVE, and average number of IVs selected as relevant (Ŝ) and relevant but invalid (Î) by TSHT. The three rows of results for each estimator correspond to sample sizes of *n* = 500, *n* = 2000 and *n* = 10, 000 respectively.

## 7 Data Application

The prevalence of type 2 diabetes mellitus is increasing across all age groups in the United States possibly as a consequence of the obesity epidemic. Many epidemiological studies have suggested that individuals with type 2 diabetes mellitus (T2D) are at higher risk of various memory impairments which are highly associated with dementia and Alzheimer’s Disease. However, such observational studies are well known to be vulnerable to confounding bias. Therefore, obtaining an unbiased estimate of the association between diabetes status and cognitive functioning is key to predicting the future health burden in the population and to evaluating the effectiveness of possible public health interventions.

In order to illustrate the proposed MR approach, we used data from the Health and Retirement Study, a cohort initiated in 1992 with repeated assessments every 2 years. We used externally validated genetic predictors of type 2 diabetes as IVs to estimate effects on memory functioning among HRS participants. The Health and Retirement Study is a well-documented nationally representative sample of persons aged 50 years or older and their spouses (Juster and Suzman,1995). Genotype data were collected on a subset of respondents in 2006 and 2008. Genotyping was completed on the Illumina Omni-2.5 chip platform and imputed using the 1000G phase 1 reference panel and filed with the Database for Genotypes and Phenotypes (dbGaP, study accession number: phs000428.v1.p1) in April 2012. Exact information on the process performed for quality control is available via Health and Retirement Study and dbGaP21 (Mailman, 2007). From the 12,123 participants for whom genotype data was available, we restricted the sample to 7,738 non-hispanic white persons with valid self-reported diabetes status at baseline and memory assessment score two years later. Self-reported diabetes in the Health and Retirement Study has been shown to have 87% sensitivity and 97% specificity for Hemoglobin A1c defined diabetes among non-Hispanic white HRS participants (White et al, 2014). Memory was assessed by immediate and delayed recall of a 10-word list plus the proxy assessments for severely impaired individuals. The validity and reliability of these measures have been documented elsewhere (Ofstedal et al. 2005; Wu et al. 2012).

Standard MR relies on the assumption that all 39 SNPs affect a person’s memory score at followup only through baseline diabetes status which is unlikely, even if all 39 SNPs only affect memory through diabetes. This is because there is likely to be a nonnegligible direct effect from one of the SNPs to diabetes incidence among persons who are diabetes-free at baseline. This would constitute a violation of the exclusion restriction and therefore would invalidate a standard MR analysis for assessing effects of baseline diabetes on memory score at follow-up. Nonetheless, although possibly positively biased under the alternative hypothesis, the two-stage regression estimator could still be interpreted as a valid test of the null hypothesis of no association between diabetes disease (whether baseline or time-updated) and memory score. It may also be true that unknown pleiotropic effects of at least one of the SNPs exists through a pathway not involving diabetes, which would constitute an even more serious violation, as it would also invalidate our MR analysis as a valid test of a causal association between diabetes and memory functioning. In light of these possible limitations a more robust MR analysis is naturally of interest.

We used GENIUS to estimate the relationship between diabetes status (coded 1 for diabetic and 0 otherwise) and memory score. As genetic instruments, we used 39 independent single nucleotide polymorphisms previously established to be significantly associated with diabetes (Morris et al 2012).

We first performed an observational analysis, which entailed fitting a linear model with memory score as outcome, diabetes status as exposure, adjusting for age at cognitive assessment and sex. Next, we implemented an MR analysis of the effects of diabetes status on cognitive score incorporating all 39 SNPs as candidate IV using TSLS, sisVIVE, adaptive LASSO, TSHT, MR Egger, and the proposed GENIUS approaches.

Participants were, on average, 68.1 years old (standard deviation [SD]=10.1 years old) at baseline and 1282 of them self-reported that they had diabetes (16.7%). The 39 SNPs jointly included in a first-stage logistic regression model to predict diabetes status explained 3.5% (Nagelkerke *R*^2^) of the variation in diabetes in the study sample, and were strongly associated as a set with the endogenous variable (Likelihood ratio test Chi-square statistic = 162 with 39 degrees of freedom, which corresponds to a significance value *<*0.001). This provides fairly compelling evidence that the IVs are not only jointly relevant but also satisfy the first stage heteroscedasticity condition required by MR GENIUS.

Table 8 shows results from both observational and IV analyses. In the observational analysis, being diabetic was associated with an average decrease of 0.04 points (s.e.=0.02) in memory score. MR GENIUS suggests a notably larger diabetes-associated decrease in average memory score equal to 0.18 points (s.e.=0.14). The efficient MR GENIUS produced a similar decrease of 0.16 points (s.e.=0.14). MR-Egger produced an estimate suggesting a protective effect of diabetes (beta=0.25, s.e.=0.35) and so did TSLS (beta=0.48, s.e.=0.22), sisVIVE (beta=0.48) and adaptive lasso (beta=0.48, s.e.=0.22) which gave the same point estimate, while TSHT (beta=0.45, s.e.=0.28) gave a slightly smaller but still protective estimate. TSLS, sisVIVE and adaptive lasso inferences coincide exactly in this application because all 39 candidate SNPs ended up being selected as”valid” by sisVIVE and adaptive lasso. In contrast, TSHT selected six candidate IVs only as both valid and relevant which were therefore used to estimate the causal effect. In conclusion, both the observational analysis and MR GENIUS found some evidence of a harmful effect of diabetes on memory score, which supports the prevailing hypothesis in the diabetes literature. In contrast, all other (robust and non-robust) MR methods suggest a protective effect of diabetes on memory, a hypothesis with little if any scientific basis in the diabetes literature.

**Table 8:** Estimation of *β*_t2d-ms_, the association between type 2 diabetes and memory score.

| | $\hat{\beta}_{\text{t2d-ms}}$ | SE | 95% CI | # of instruments<br>selected as invalid |
| --- | --- | --- | --- | --- |
| Observational analysis |  |  |  |  |
|  | -0.04 | 0.02 | (-0.08, 0.001) | - |
| IV analyses |  |  |  |  |
| MR GENIUS | -0.18 | 0.14 | (-0.45, 0.08) | - |
| Efficient MR GENIUS | -0.16 | 0.14 | (-0.43, 0.11) | - |
| MR-Egger | 0.25 | 0.35 | (-0.43, 0.93) | - |
| sisVIVE | 0.48 | - | - | 0 |
| TSLS | 0.48 | 0.22 | ( 0.05, 0.90) | - |
| Adaptive Lasso | 0.48 | - | - | 0 |
| Post-adaptive Lasso | 0.48 | 0.22 | ( 0.05, 0.90) | 0 |
| TSHT | 0.45 | 0.28 | (-0.10, 1.00) | 0 (out of 6<br>selected as relevant) |

## 8 Concluding Remarks

As MR gains popularity as a promising strategy to address confounding bias in observational studies, there clearly also is a growing need for robust MR methodology that relax the standard IV assumptions. Although a variety of methods have recently been proposed, we have argued that MR GENIUS stands out as an effective approach with clear advantages over other existing methods. First, the approach bypasses the overly stringent orthogonality condition of MR-Egger. Furthermore, whereas existing methods are technically only consistent either as the number of candidate IVs goes to infinity (MR Egger), or as a majority (adaptive lasso) or a plurality (TSTH) of IVs are valid, MR GENIUS is guaranteed to be consistent without even one valid IV. Furthermore, as we have shown, MR GENIUS equally applies with continuous or binary outcome, continous or binary exposure and IV, multiple IVs, auxiliary pre-IV covariates and under both prospective and retrospective sampling designs. Finally, whereas adaptive lasso and TSTH require one or more model selection steps therefore compromising inferences that are uniform over the entire model of interest, MR GENIUS does not involve model selection, therefore bypassing this difficulty.

MR GENIUS also stands out from other methods because it does not require modeling the effects of invalid IVs on *Y* for consistent estimation of the effect of exposure, therefore allowing main effects and interactions among components of *G* to remain unrestricted in the outcome model. In the event of an interaction between *A* and *G,* such that equation (1) does not hold, it is straightforward to show that MR GENUIS estimates a certain weighted average of the causal effect. For instance, in the case of binary *A, µ* = *E* (*β* (*G*) *ϖ* (*G*)) where *β* (*G*) = *E* (*Y |A* = 1*, G*) *- E* (*Y |A* = 0*, G*) is the causal effect within levels of *G* and *ϖ*(*G*) = (*G - E*(*G*)) *var*(*A|G*) *× E{*(*G - E*(*G*)) *var*(*A|G*)*}*^*-*1^; or equivalently *µ* = *β* (0) + (*β* (1) *- β* (0)) *× var*(*A|G* = 1)*var*(*G*) *× E{*(*G - E*(*G*)) *var*(*A|G*)*}*^*-*1^. Therefore, MR GENUIS is guaranteed to be consistent under the null hypothesis of no conditional effect of exposure within levels of *G* provided there is no interaction between *U* and *G* in the outcome model. An R package which implements MR GENIUS is available at github.com/bluosun/MR-GENIUS.

In closing, we acknowledge certain limitations of MR GENIUS. First, the approach may be vulnerable to weak IV bias which may occur if *var*(*A|G*) is weakly dependent on *G*, a possibility that was largely ruled out in this paper. MR GENIUS is also currently not designed to handle high dimensional IVs (where the number of IVs may exceed sample size). We plan to further develop MR GENIUS to address all of these remaining challenges in future work.

## 9 Acknowledgment

Eric Tchetgen Tchetgen’s work is funded by NIH grant R01AI104459. The Health and Retirement Study genetic data are sponsored by the National Institute on Aging (grant numbers U01AG009740, RC2AG036495, and RC4AG039029) and was conducted by the University of Michigan. The authors thank Frank Windmeijer for valuable discussions.

## Appendix

### A1. Proof of Lemma 5

**Proof.** We first note that for any additive function *t*(*A, G*) = *t*_1_(*A*) + *t*_2_(*G*),

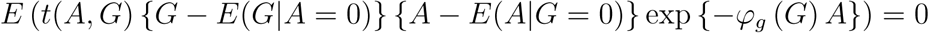

because

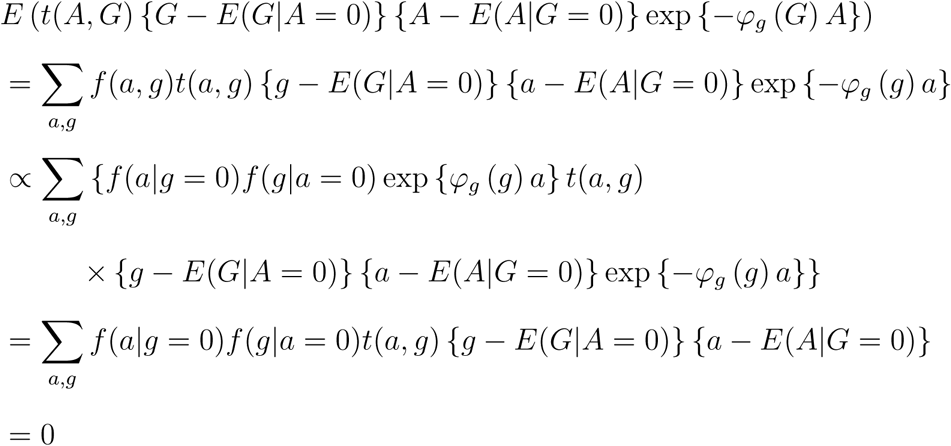

where we used the fact that

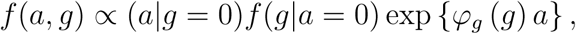

see for example Tchetgen Tchetgen et al (2009). It is straightforward to verify that the

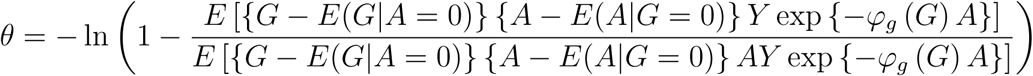

Next

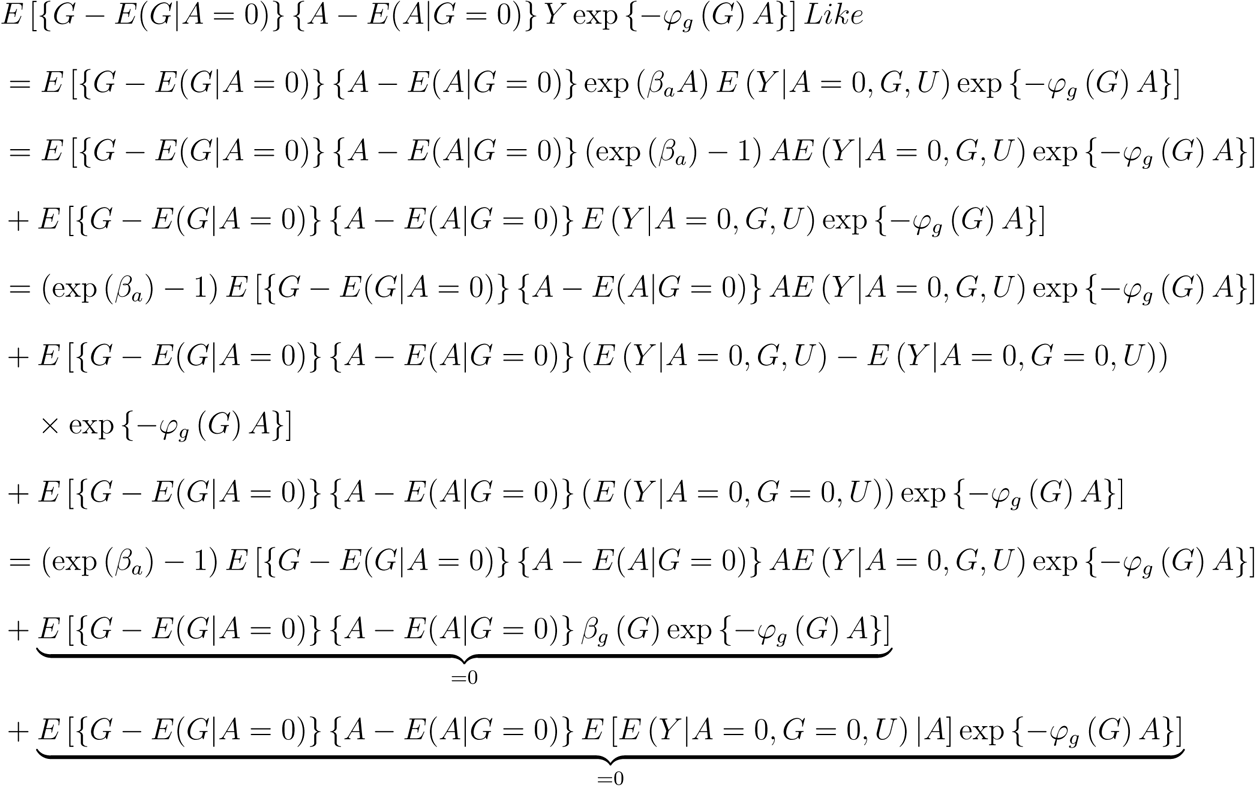

Likewise

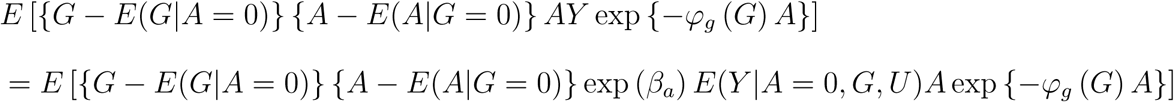

Therefore

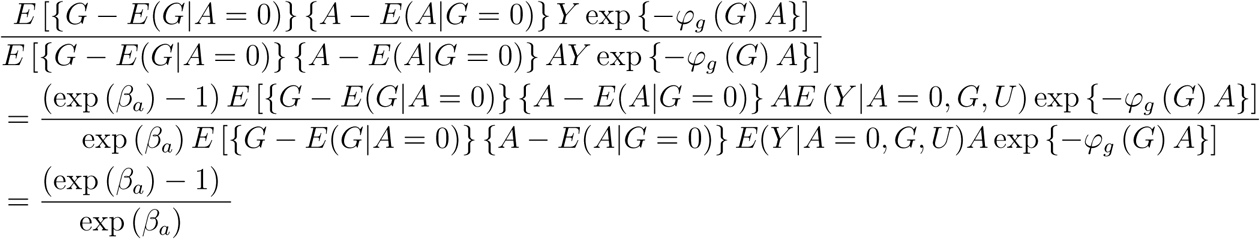

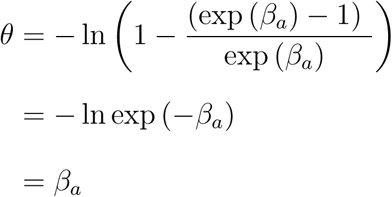

provided that

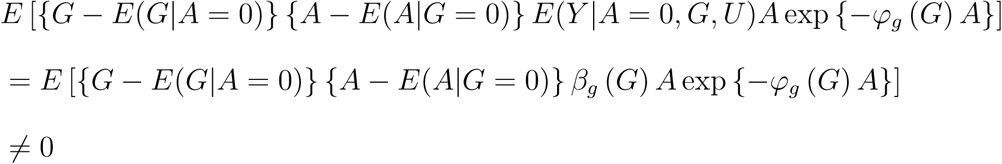

which holds by assumption because *γ*_*ag*_(*g*) = (exp (*β*_*a*_) *-* 1) *β*_*g*_ (*G*).

### A2. Variance estimation with single IV

The estimating equation in (6) involves the estimated nuisance parameters 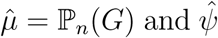 of the model *E*(*A|G*; *ψ*). To account for the effect of nuisance parameter estimation on the subsequent estimation of *β*_*a*_, the empirical moment conditions are stacked to form

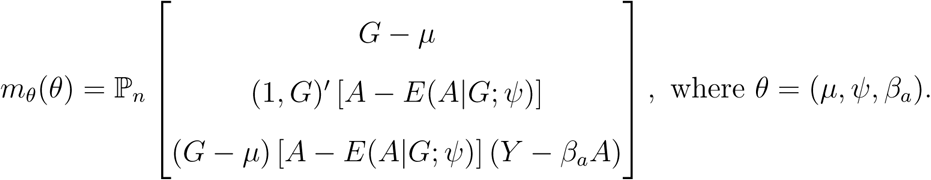

The estimation procedure satisfies the joint conditions 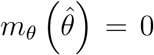. Without loss of generality, we specify [*A - E*(*A|G*; *ψ*_0_)] as a main effects model with intercept. Assume standard regularity conditions and expand 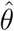 around the true parameter value *θ*_0_ yields

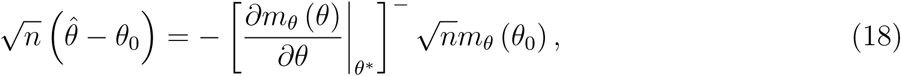

where *θ\** is intermediate in value between 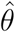 and *θ*_0_. It follows that

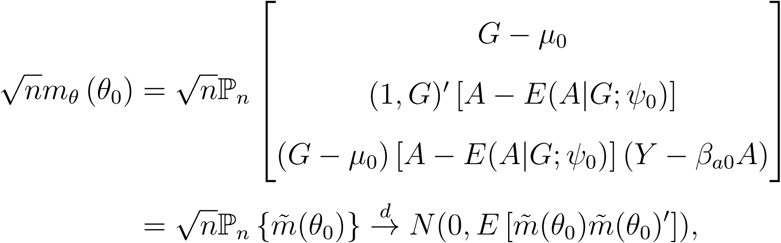

while for the”bread” matrix

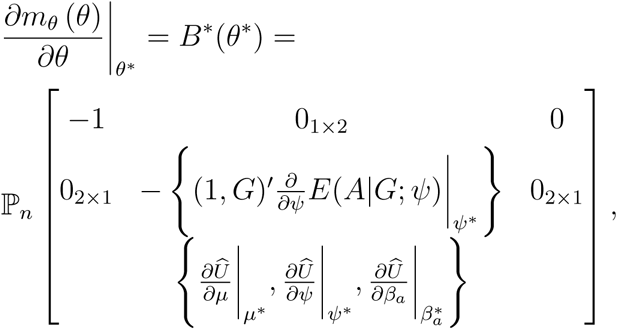

where

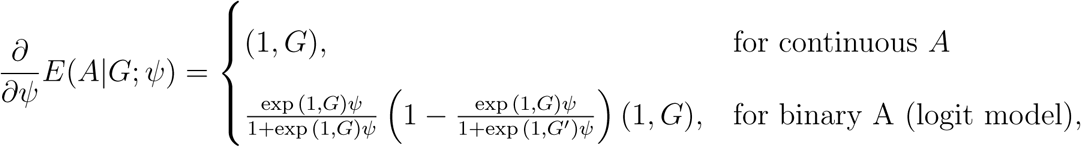

and

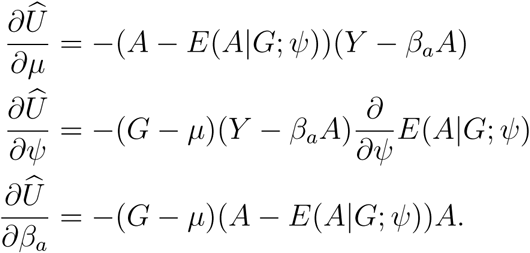

Assume that the matrix *B*(*θ*_0_) is non-singular, where the entries in *B*(*θ*_0_) are the expected values of the sample averages in *B\**(*θ\**), evaluated at *θ*_0_. Then 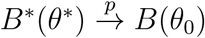, and

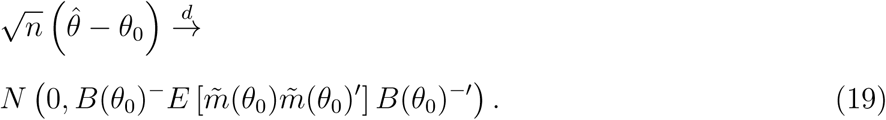

Replacing the expected values in (19) with sample averages evaluated at 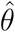 yields a consistent estimator of the asymptotic covariance matrix. For inference about *β*_*a*_, one may report its Wald-type 95% confidence interval constructed with the corresponding component of the estimated covariance matrix for 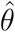.

### A3. Variance estimation with multiple IVs

Let 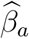 be the solution to (12) with optimal weight 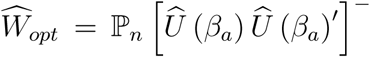 where *T*^−^ denotes the generalized inverse of matrix *T*. The empirical moment conditions 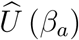 in (12) involves the first stage estimates 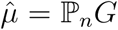 as well as 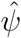of the model *E*(*A|G*; *ψ*), which effects need to be accounted for in the subsequent estimation of *β*_*a*_. Without loss of generality, we specify [*A - E*(*A|G*; *ψ*_0_)] as a main effects model with intercept. If there are *k* IVs, let

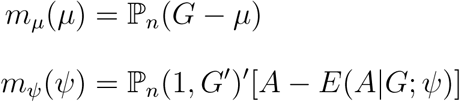

be the *k* and (*k* + 1) empirical moment conditions of obtaining 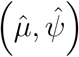 respectively. For iterated or continuously updated GMM procedures in which *β*_*a*_ is estimated simultaneously with the optimal weight, the first order condition of (12) is

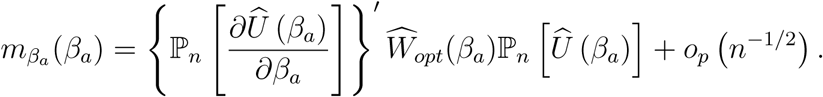

The two-stage procedure solution satisfies the joint moment conditions

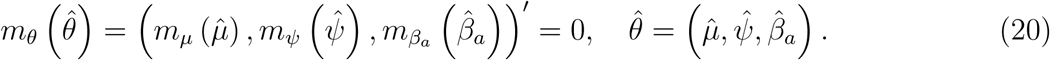

Assume standard regularity conditions and expand 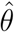 around the true parameter value *θ*_0_ yields

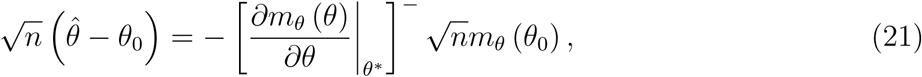

where *θ\** is intermediate in value between *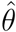* and *θ*_0_. Consider

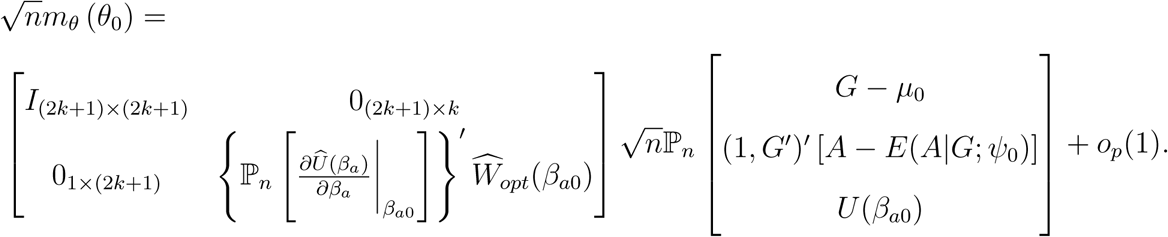

Let

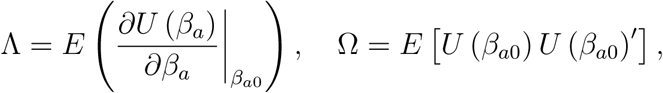

so that

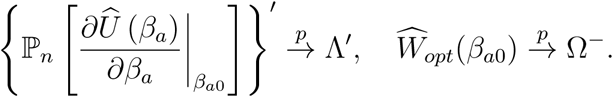

Then

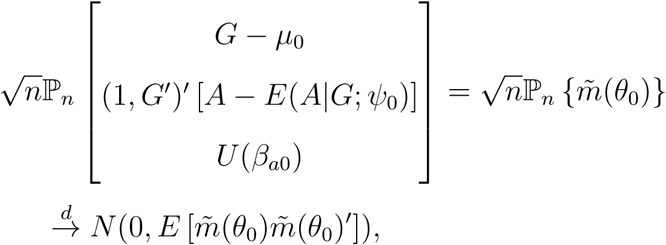

and by Slutsky’s theorem

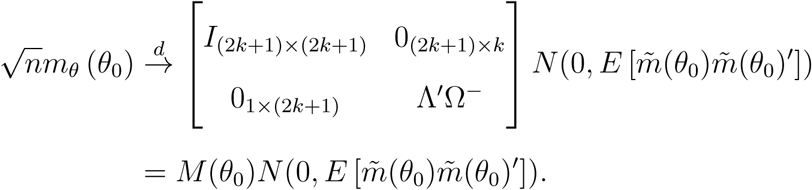

Next consider the”bread” matrix

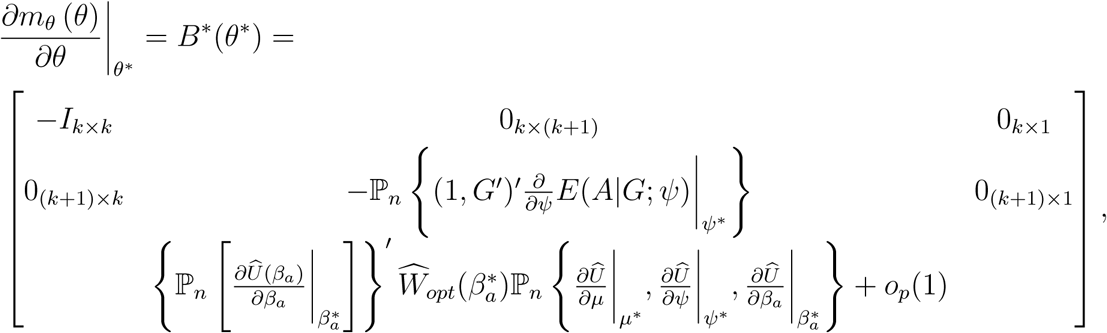

where

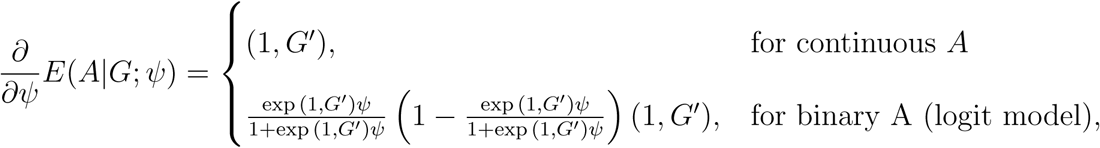

and

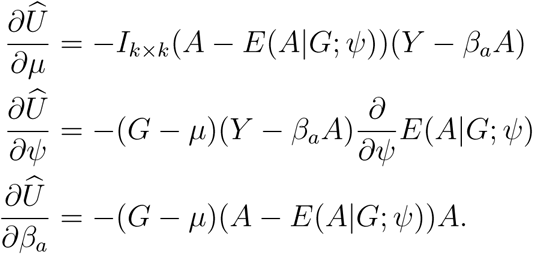

Assume that the matrix *B*(*θ*_0_) is non-singular, where the entries in *B*(*θ*_0_) are the expected values of the sample averages in *B\**(*θ\**), evaluated at *θ*_0_. Then 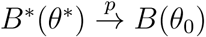, and

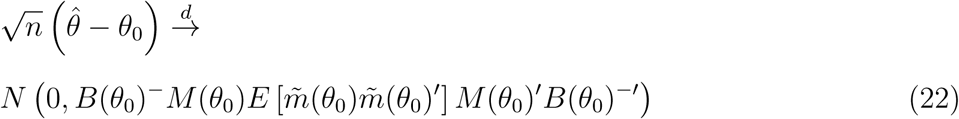

In practice, replacing the expected values in (22) with sample averages evaluated at 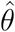 yields a consistent estimator of the asymptotic covariance matrix. In addition, centering the IV moment conditions Û (*β*_*a*_) when estimating the covariance matrix 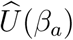 may improve finite sample inference. For inference about *β*_*a*_, one may report its Wald-type 95% confidence interval constructed with the corresponding component of the estimated covariance matrix for 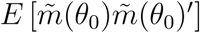. The above variance estimation framework can accommodate baseline covariates *C* by stacking the moment conditions for Ê(*G|C*) and *Ê*(*A|G, C*) instead, as described in estimating equation (10)

